# Tricuspid valve regurgitation accelerates heart failure via a cardio-intestinal innate immune circuit

**DOI:** 10.64898/2026.07.07.736969

**Authors:** Florian Sicklinger, Tobias Thiemann, Sarah Rupprecht, Lisa Quadt, Junedh M. Amrute, Jan Zuchgan, Jakob C. Voran, George Markousis-Mavrogenis, Alejandra Isasi Nalvarte, Laura M. Wienecke, Niklas Hartmann, Stephan Erbe, Isabel A. Hoerbrand, Martin J. Kraus, Martin Gruber, Robert Bibernell, Simone Martini, Lucia S. Kilian, Hauke Hund, Jes-Niels Boeckel, Matthias Mack, Adriaan A. Voors, Peter van der Meer, Derk Frank, Norbert Frey, Kory J. Lavine, Mathias Konstandin, Florian Leuschner

## Abstract

Activation of the immune system impacts the progression of heart failure (HF), but the underlying mechanisms remain incompletely understood. Here, we identify a cardio-intestinal innate immune axis that links systemic venous congestion to myocardial inflammation, fibrosis, and functional decline. Using single-cell and single-nucleus transcriptomic profiling in patients and mice with tricuspid regurgitation (TR), we demonstrate that TR disrupts intestinal barrier integrity and elicits expansion of circulating monocytes which in turn orchestrate pathological crosstalk between the right and left heart. Monocyte-derived Interleukin-6 (IL-6) emerged as a key mediator of TR-driven myocardial fibrosis and dysfunction. Blockade of IL-6 attenuated cardiac fibrosis and improved cardiac function. In patients, catheter-based repair of TR resulted in reduced IL-6 levels. Together, these findings establish cardio-intestinal innate immunity as a mechanism linking altered hemodynamics to left ventricular remodeling and nominate TR patients as a selective target population for IL-6–directed therapy in HF.

**One Sentence Summary:** This work mechanistically resolves the heart-gut axis in tricuspid valve regurgitation, and its impact on heart failure progression as mediated by Interleukin-6.

## INTRODUCTION

The mammalian heart consists of two anatomically and functionally distinct ventricles that operate in series to sustain systemic and pulmonary circulation. Although heart failure (HF) is often viewed as a disease of the left ventricle (LV), both ventricles function as an integrated unit, and right ventricular performance is increasingly recognized as a critical determinant of overall cardiac function (*1*). Functional impairment of the right ventricle (RV) affects up to 50% of all patients with left-sided HF (*2, 3*). Although RV dysfunction strongly predicts adverse outcomes, it remains unclear whether it is a mere bystander of left-sided heart disease or whether it actively drives left ventricular remodeling and dysfunction (*4, 5*). Mechanistic pathways underlying this biventricular interaction are poorly understood, and no targeted therapies exist for RV failure (*6, 7*).

Tricuspid valve regurgitation (TR) is a common manifestation of acute and chronic RV failure and remodeling. Affecting more than 4 million individuals in the United States and Europe, TR is associated with substantial morbidity and mortality (*8–10*). The most prevalent form, functional (secondary) TR, arises from tricuspid annular dilation and architectural changes in the subvalvular apparatus (*1, 2*). Consequently, regurgitant flow from the RV into the right atrium during each systole contributes to chronic venous congestion, affecting multiple organ systems including the splanchnic circulation. Venous congestion is a well-recognized feature of disease progression in both acute and chronic HF, with links to end-organ dysfunction, immune activation, and systemic inflammation (*11*). Yet whether congestion simply reflects advanced disease or actively contributes to HF progression remains unresolved. Addressing this question has been challenging because clinical studies are largely observational and associative (*12*). Further, experimental models capable of isolating the effects of chronic venous congestion in HF have been lacking. Therefore, defining how TR-associated venous congestion influences cardiac remodeling and biventricular communication is a critical unmet challenge.

Here, we explored systemic and heart-specific molecular alterations in patients with severe TR and matched controls. TR was associated with increased left ventricular fibrosis and a robust expansion of pro-inflammatory monocytes. To determine the mechanistic processes of TR-induced inflammation and remodeling, we developed a novel mouse model enabling induction of TR, either in isolation or in the context of pre-existing left ventricular dysfunction. In mice, TR accelerated left ventricular failure and adverse remodeling. Single-nucleus transcriptomics combined with monocyte depletion and adoptive transfer experiments identified circulating monocyte–derived Interleukin-6 (IL-6) as a key mediator driving left ventricular remodeling and fibrosis. Therapeutic blockade of IL-6 in HF with concomitant TR attenuated cardiac fibrosis and improved outcomes in vivo. Collectively, these findings establish TR-driven inflammation as a causal link between right heart pathology and adverse left ventricular remodeling. We propose immune-targeted therapies as potential disease-modifying interventions for TR-associated HF.

## RESULTS

### TR is linked to LV dysfunction and fibrosis in patients

To assess the impact of TR on left ventricular remodeling in patients, we first conducted a retrospective matched cohort study within a clinical population of 1,299 patients with left ventricular HF defined by a reduced or mildly reduced ejection fraction (EF <50%; encompassing HFrEF and HFmrEF). We identified 92 cases with severe TR and matched each to four controls based on age, sex and LV ejection fraction (Fig. 1A, table S1). Consistent with prior studies (*13*), TR was associated with increased all-cause mortality over 5 years (Fig. 1B) and with worse LV ejection fraction after 1-year follow-up (Fig. 1C), suggesting a causal contribution to adverse cardiac remodeling. To define the molecular correlates of this observation, we analyzed single-nucleus RNA sequencing (snRNA-seq) data from left ventricular tissue of patients with HFrEF (*14*), stratified by the presence or absence of concomitant TR (Fig. 1D). A total of 66,669 nuclei from six TR patients and seven control patients were analyzed and partitioned into clusters using established lineage markers (Fig. 1E, fig. S1A). Left ventricular tissue from patients with TR exhibited a marked relative expansion of fibroblast populations compared with non-TR controls (Fig. 1F), while the proportions of all other major cardiac cell types remained comparable between groups (fig. S1B-G). This expansion was predominantly driven by an increase in a Serpine1+ fibroblast subcluster (Fig. 1G, fig. S1H-I), a population previously implicated in pro-fibrotic extracellular matrix remodeling (*14*). Differential gene expression and gene set enrichment analysis of the expanded fibroblast pool revealed enrichment of gene programs related to extracellular matrix organization and inflammatory signaling (Fig. 1H, fig. S1J), including cellular responses to IL-6. Thus, left ventricular fibroblasts undergo substantial pro-fibrotic and inflammatory remodeling during their expansion in patients with TR.

**Fig. 1.**
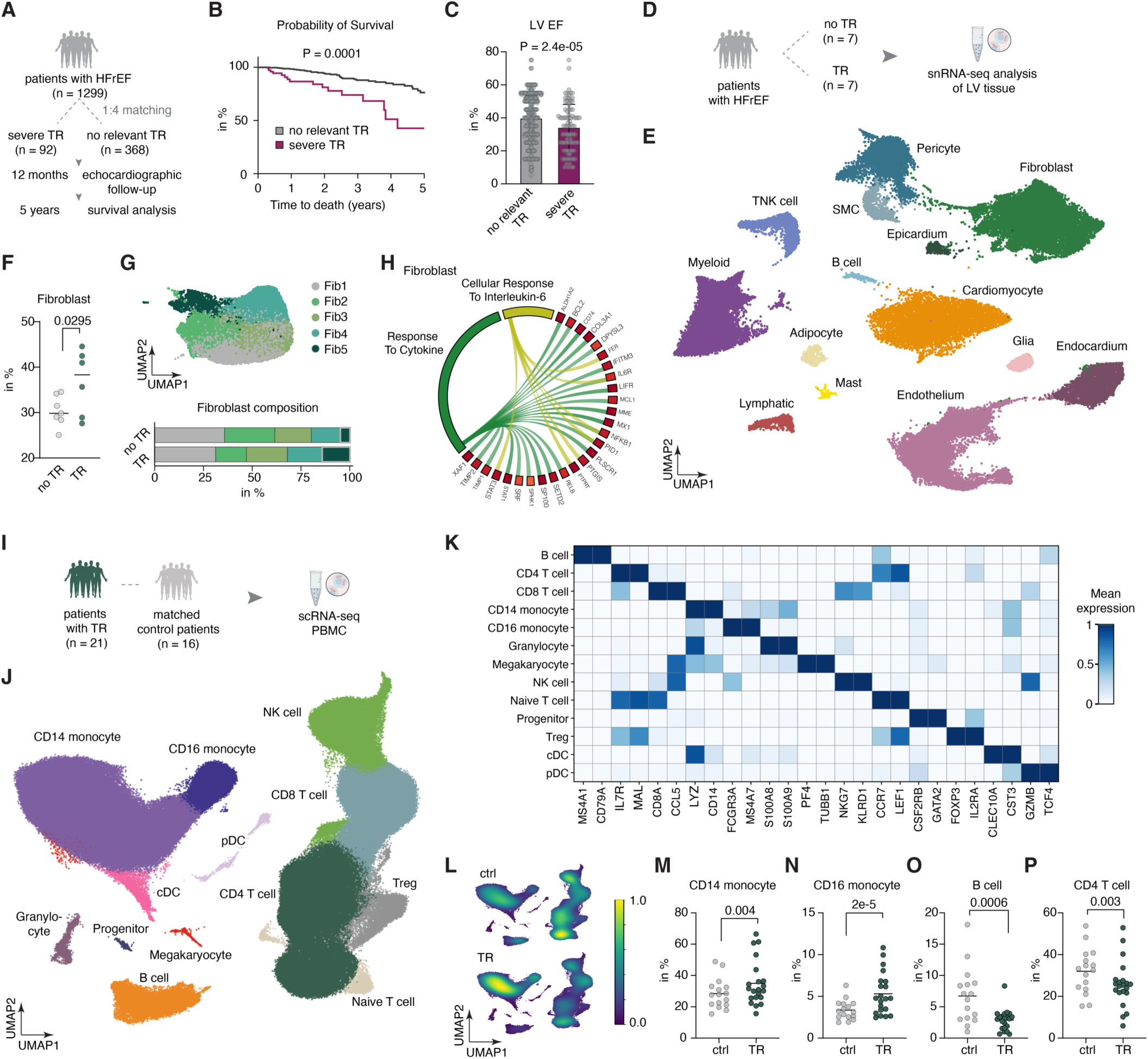
TR is associated with adverse left ventricular remodeling and systemic monocyte activation in patients. (**A**) Schematic of the retrospective matched cohort study design. Patients with severe TR (n=92) were matched 1:4 to controls by age, sex, and LV EF within a HF cohort (EF <50%; n=1,299). (**B**) Kaplan-Meier curves for all-cause mortality over 5 years (multivariate-regression analysis, *P*-value shown for group effect). (**C**) Quantification of LV EF at 12-month follow-up in TR (magenta) versus control (gray) patients (multivariate-regression analysis, *P*-value shown for group effect, mean ± SD with individual data points). (**D**) Schematic of the snRNA-seq study design. LV tissue from TR (n=6) and control (n=7) HFrEF patients was profiled. (**E**) UMAP embedding of 66,669 single nuclei from LV tissue. SMC, smooth muscle cell. (**F**) Fibroblast proportion (% of total cells) per sample in TR versus control patients. Data show mean with individual data points. *P*-values were computed using weighted least squares (WLS) regression. (**G**) UMAP of cardiac fibroblast subclusters (top) and proportional composition per condition (bottom). (**H**) Chord diagram depicting identified gene sets enriched in TR fibroblasts and their contributing upregulated genes. (**I**) Schematic of the scRNA-seq PBMC study design. PBMCs from 21 TR patients and 16 matched controls were profiled. (**J**) UMAP embedding of 345,821 single cells colored by annotated cell type. NK, natural killer; cDC, conventional dendritic cell; pDC, plasmacytoid dendritic cell; Treg, regulatory T cell. (**K**) Heatmap of marker gene expression used for cell type assignment. (**L**) Density plots of cell distribution across the UMAP for TR patients (top) and controls (bottom); color scale from low (blue) to high (yellow) density. (**M-P**) Cell type frequency (% of total PBMCs) for CD14+ monocytes (M), CD16+ monocytes (N), B cells (O), and CD4+ T cells (P). Data show means with individual data points. *P*-values were calculated using WLS regression with Benjamini-Hochberg correction.

### Peripheral monocytes expand in human TR

The enrichment of inflammatory signaling pathways in left ventricular fibroblasts suggested that TR-associated remodeling may be coupled to systemic immune activation. To validate this, we collected peripheral blood mononuclear cells (PBMCs) from 16 control participants and 21 patients with severe TR (Fig. 1I, table S2) and performed single-cell RNA sequencing (scRNA-seq) on the Parse platform (*15*). Following quality control, doublet removal, data integration and clustering, we identified 345,821 single cells partitioned into 13 major populations, including monocytes, lymphoid cells, natural killer cells, granulocytes, progenitor cells, and megakaryocytes (Fig. 1J–K). In patients with TR, both CD14+ classical and CD16+ non-classical monocyte populations were significantly expanded (Fig. 1L–N), whereas B cells and CD4+ T cells were decreased (Fig. 1O-P). Among less abundant cell populations, regulatory T cells, naive T cells, and granulocytes also showed significant compositional shifts (fig. S2A–I). Differential expression analysis revealed widespread transcriptional alterations within monocytes. Monocytes from patients with TR exhibited a pro-inflammatory phenotype, characterized by significant upregulation of genes involved in macrophage activation and response to biotic stimulus (fig. S2J–K). Expansion of circulating monocytes due to TR could be validated in a separate cohort of patients with HF with reduced ejection fraction (*16*) (fig. S2L–M). Taken together, these findings indicate that TR is associated with chronic systemic immune activation marked by expansion and inflammatory reprogramming of circulating monocytes.

### A mouse model of TR

To dissect immune pathways involved in human TR mechanistically, we developed a mouse model of TR (Fig 2A). We used a trans-LV access to sever the tricuspid valve using 27-gauge micro scissors under echocardiographic guidance (fig. S3A-B, movie S1). This minimally invasive approach yielded reproducible induction of regurgitant jet at the tricuspid valve (Fig. 2B, fig. S3C). Ventricular septal defects after the procedure due to the access route could be excluded using color Doppler imaging (fig. S3D). The induction of TR elicited right ventricular and right atrial enlargement with significant dilation of the inferior vena cava (IVC), commonly found in patients with TR (Fig. 2C–F, fig. S4A–B). Invasive hemodynamic measurements revealed right ventricular volume overload without significant increase in right ventricular end-systolic pressure (Fig. 2G–H, fig. S4C–F). In TR hearts, the RV exhibited hypertrophy and dilation (fig. S4G–H), accompanied by increased cardiomyocyte size, interstitial fibrosis, and macrophage accumulation (fig. S4I–K). Left ventricular function after isolated TR induction was not significantly changed (fig. S4L–M). Furthermore, we found no differences in LV cardiomyocyte size or fibrotic tissue content (fig. S4N–O). However, immunofluorescence stainings revealed a higher abundance of CD68+ cardiac macrophages in left ventricular tissue which was even more prevalent in the interventricular septum (fig. S4P–R). Of note, we did not observe any mortality peri-procedurally nor in a follow-up period of 12 weeks after TR induction.

**Fig. 2.**
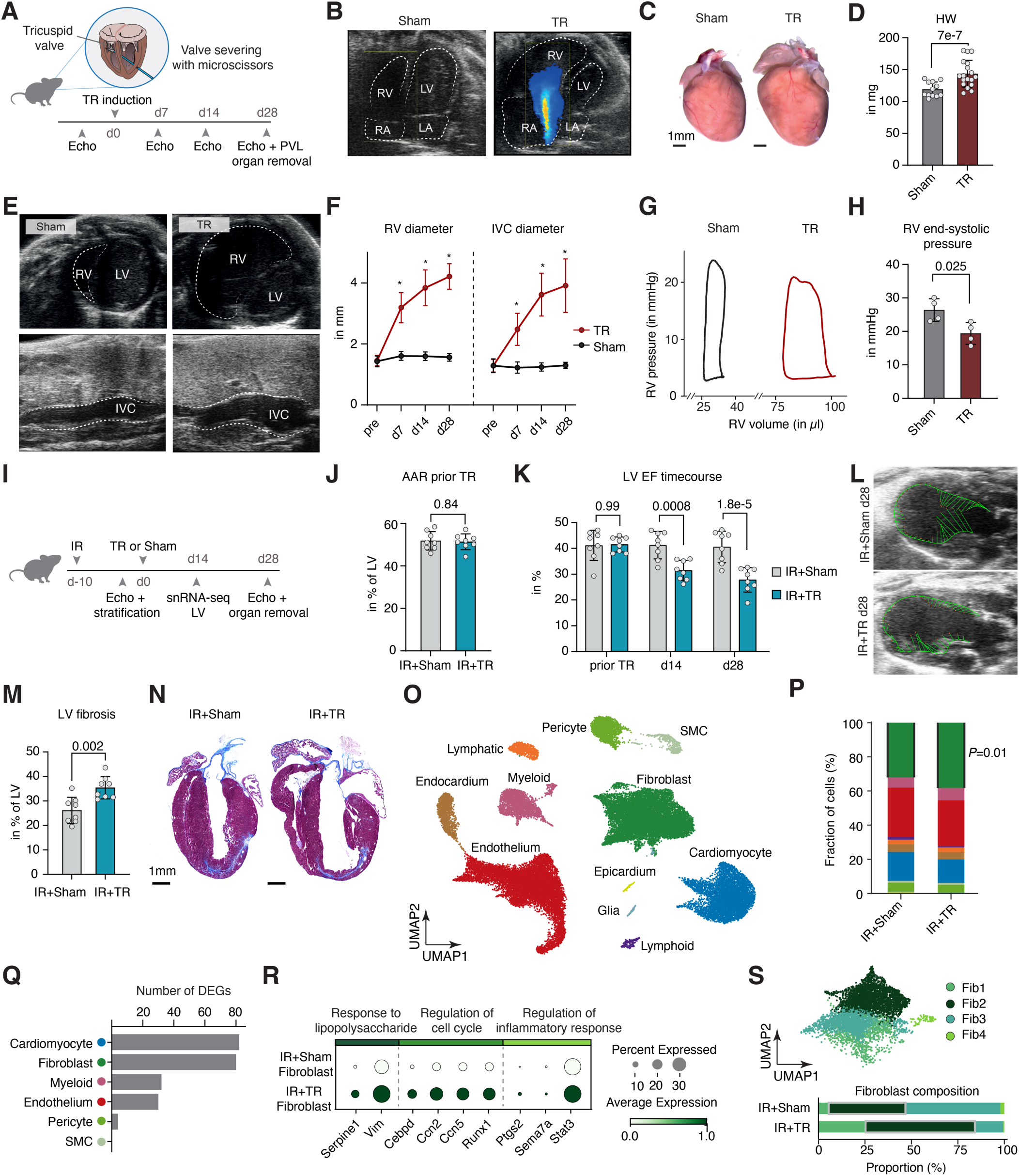
TR in mice induces right ventricular remodeling and accelerates left ventricular failure in chronic ischemic cardiomyopathy. (**A**) Schematic and experimental timeline for echocardiography-guided TR induction via trans-LV microsurgical tricuspid valve severing in mice. (**B**) Representative Color Doppler echocardiographic images from sham and TR mice 28 days post-intervention. RV, right ventricle; LV, left ventricle; RA, right atrium; LA, left atrium. (**C**) Representative gross cardiac specimens from sham and TR mice at 4 weeks. Scale bar, 1mm. (**D**) Heart weight at d28 post-intervention (n = 12-18, unpaired t-test, mean ± SD). (**E**) Representative echocardiographic parasternal short-axis (top) and subcostal IVC (bottom) images at d28. IVC, inferior vena cava. (**F**) Serial echocardiographic quantification of RV diameter (left) and IVC diameter (right) over time (n = 8, two-way ANOVA with Sidak’s multiple comparison, mean ± SD). \**P* < 0.05. (**G**) Representative invasive RV pressure-volume loops at 4 weeks. (**H**) RV end-systolic pressure at d28 (n = 4, unpaired t-test, mean ± SD). (**I**) Experimental timeline for superimposed TR in chronic ischemic cardiomyopathy: TR or sham intervention was performed 10 days after ischemia-reperfusion (IR) injury, with terminal assessment at d28 post-TR. (**J**) Area at risk (AAR) quantified by echocardiography prior to TR induction to stratify groups (n = 8 per group, unpaired t-test, mean ± SD). (**K**) Serial LV EF at indicated timepoints across both groups (n = 8, two-way ANOVA with Sidak’s multiple comparison, mean ± SD). (**L**) Representative B-mode echocardiographic images at d28 in IR+sham (top) and IR+TR (bottom) mice with LV endocardial movement tracings (green). (**M**) LV fibrotic area as a percentage of AAR at d28 (n = 8, unpaired t-test, mean ± SD). (**N**) Representative Masson’s trichrome stained long-axis sections at d28. Scale bar, 1mm. (**O**) UMAP of snRNA-seq data from LV nuclei across 4 conditions (baseline+sham, baseline+TR, IR+sham, IR+TR) collected 14 days after TR or sham intervention. SMC, smooth muscle cell. (**P**) Stacked bar plots showing proportional composition of major cardiac cell types per condition based on snRNA-seq (WLS regression, Benjamini–Hochberg correction, *P*-value shown for Fibroblast). (**Q**) Number of differentially expressed genes (DEGs) per cell population between IR+TR and IR+sham conditions, identified by pseudobulk DESeq2 (n = 3 per group, Wald test, Benjamini–Hochberg, *P*adj. < 0.05). (**R**) Dot plot of DEG expression in fibroblasts from IR+sham (top) and IR+TR (bottom) conditions, grouped by enriched GO terms. (**S**) UMAP of cardiac fibroblast subclusters (top) and corresponding proportional composition across conditions (bottom).

### TR accelerates left ventricular HF

We next sought to validate the translational relevance of these findings in the context of left-sided HF. Because TR frequently coexists with left ventricular dysfunction, we employed a chronic HF model induced by myocardial ischemia-reperfusion (IR) injury (*17*), the most common cause of HF with reduced ejection fraction (*18*). To minimize confounding effects of acute post-infarction remodeling, TR or sham intervention was performed 10 days after IR (Fig 2I), a time-point at which infarct inflammation is resolved (*19*). Following stratification by scar size and LV ejection fraction (Fig. 2J–K), mice were subjected to either TR induction or sham surgery. Notably, superimposed TR significantly exacerbated pre-existing left-sided HF, as evidenced by a further reduction in LV systolic function (Fig. 2K–L). Consistently, mice with concomitant TR exhibited more pronounced left ventricular dilation and increased heart-to-body weight ratios (fig. S5A-B), indicative of aggravated cardiac remodeling. Since ischemic HF is characterized by progressive myocardial fibrosis, we next quantified fibrotic remodeling. One month after TR induction, mice with combined IR and TR showed significantly increased left ventricular fibrosis compared with IR controls (Fig. 2M–N, fig. S5C–D). To define the molecular basis of TR-induced worsening of left-sided HF, we performed snRNA-seq of left ventricular tissue from four experimental groups (baseline+sham, baseline+TR, IR+sham, and IR+TR) 14 days after TR or sham intervention. After quality control and unsupervised clustering, 29,070 nuclei were retained, enabling annotation of all major cardiac cell types (Fig. 2O, fig. S5E). Comparison across groups revealed condition-specific compositional differences (fig. S5F). Notably, superimposed TR induced an expansion of fibroblasts in the context of pre-existing HF (Fig. 2P). Differential gene expression analysis revealed pronounced transcriptional changes, particularly in cardiomyocytes and fibroblasts, whereas other cell populations were comparatively less affected (Fig. 2Q). Fibroblasts from hearts with superimposed TR exhibited a transcriptomic signature consistent with activation (Fig. 2R), including upregulation of genes involved in cell cycle regulation (e.g., Ccn2, Ccn5, Runx1), inflammatory signaling (e.g., Ptgs2, Sema7a, Stat3), and responses to lipopolysaccharide (LPS, e.g., Serpine1, Vim). Subclustering further identified a relative expansion of a pro-fibrotic Postn+ Serpine1+ fibroblast population, consistent with an activated, matrix-producing phenotype (Fig. 2S, fig. S5G–H).

### Chronic TR spurs IL-6 release by circulating monocytes

Given the inflammatory fibroblast phenotype in left ventricular tissue of both patients and mice with TR, and the pro-inflammatory reprogramming of circulating monocytes in patients, we next examined peripheral immune cell alterations in the murine TR model. Consistent with the findings in patients, experimental TR induced an expansion of peripheral Ly6C-high monocytes during the chronic phase (Fig. 3A), whereas other leukocyte populations remained largely unchanged (fig. S6A–D). Splenic Ly6C-high monocytes were similarly expanded (fig. S6E–I), while bone marrow Ly6C-high monocytes were selectively reduced. In contrast, hematopoietic stem and progenitor cells remained unaltered (fig. S6J–R), suggesting enhanced peripheral mobilization rather than augmented de novo hematopoiesis. Monocytes are potent activators of fibroblasts through secreted mediators (*20*). Since cardiac monocyte numbers were not proportionally increased in our snRNA-seq analysis of left ventricular tissue, we hypothesized that soluble mediators — rather than direct cellular infiltration — might underlie the observed fibroblast response. We therefore measured 48 cytokines in serum of mice 4 weeks after TR or sham induction using proximity extension assay technology (Olink). We observed pronounced alterations in circulating cytokines, with IL-6 emerging as the dominant systemically regulated factor (Fig. 3B–C). Importantly, serum IL-6 levels correlated with the degree of venous congestion, as assessed by IVC diameter (Fig. 3D). Having established that TR drives concomitant expansion of circulating monocytes and systemic IL-6 elevation, we next sought to determine whether these two observations are mechanistically linked. IL-6 is a key cytokine produced by myeloid cells under inflammatory conditions (*21*), yet other cell types and organs including the liver represent established sources under certain pathological conditions (*22, 23*). To distinguish between these possibilities, we quantified IL-6 protein expression across tissues and found predominant upregulation in serum rather than in solid organs (fig. S7A). Consistent with this, hepatic mRNA levels of IL-6 were not significantly elevated during chronic TR (fig. S7B). In contrast, flow cytometric analysis revealed a significant increase in IL-6–producing monocytes in blood of TR mice (Fig. 3E–F, fig. S7C). Antibody-mediated depletion of circulating monocytes markedly reduced serum IL-6 levels (Fig. 3G, fig. S7D–E), establishing these cells as the principal systemic source in this context (*24*).

**Fig. 3.**
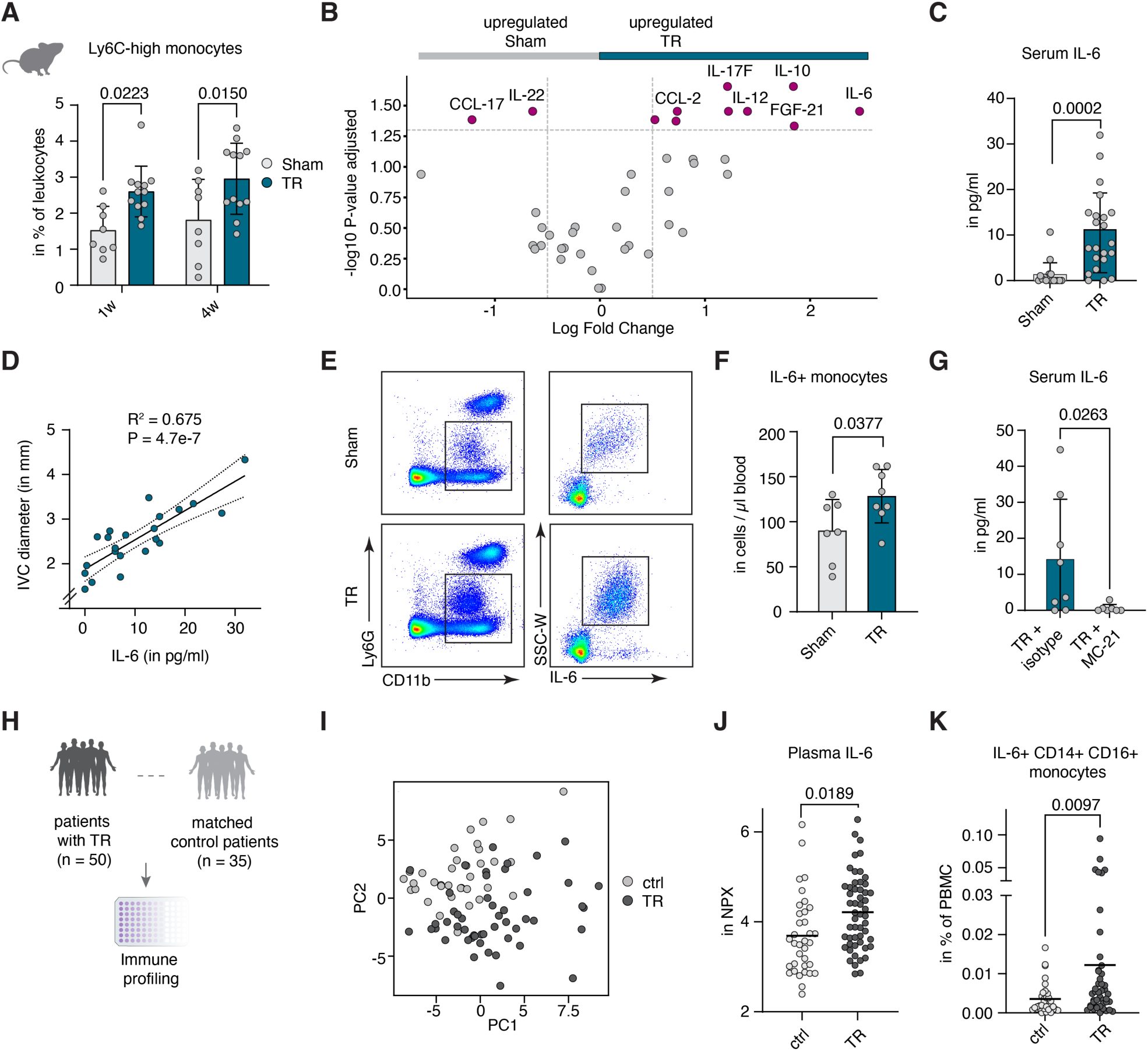
Expansion of IL-6–producing monocytes and systemic IL-6 elevation in TR. (**A**) Frequency of Ly6C-high monocytes (% of CD45+ cells) in peripheral blood at 1 and 4 weeks post-TR or sham induction (n = 8-12, two-way ANOVA with Sidak’s multiple comparison, mean ± SD). (**B**) Volcano plot of serum cytokines differentially regulated between TR and control mice based on Olink Target 48 Mouse Cytokine; colored points indicate *P*adj. < 0.05 (n = 3, unpaired t-test, Benjamini–Hochberg correction). (**C**) Serum IL-6 concentrations 4 weeks after TR or sham induction (n = 17-22, Mann-Whitney test, mean ± SD). (**D**) Correlation between serum IL-6 and IVC diameter as a measure of venous congestion severity in mice with TR on d28; R² and *P*-value by linear regression (n = 22). (**E**) Representative flow cytometry gating for IL-6+ monocytes in blood of TR and sham mice at day 28. (**F**) Quantification of IL-6+ monocytes in peripheral blood 4 weeks post-TR or sham (n = 7-8, unpaired t-test, mean ± SD). (**G**) Serum IL-6 levels following antibody-mediated monocyte depletion in TR mice (n = 7-8, Mann-Whitney test, mean ± SD). (**H**) Schematic of Olink proteomics profiling of plasma from 50 TR patients and 35 matched controls. (**I**) Principal component analysis of 92 circulating proteins from all profiled patients; TR patients (dark grey) and controls (light grey). (**J**) Plasma IL-6 (in Olink NPX units) in TR patients and matched controls (n = 35-50, unpaired t-test, Benjamini–Hochberg correction, mean with individual data points). (**K**) Frequency of IL-6+ CD14+ CD16+ monocytes in PBMCs from TR patients and matched controls measured by flow cytometry (n = 50 TR, n = 35 controls, Mann Whitney test, mean with individual data points).

To extend these findings to humans, we also profiled patients with clinically relevant TR and matched controls using the Olink proximity extension assay technology (Fig. 3H, table S3). Principal component analysis of circulating proteins revealed a separation between patients with TR and matched controls (Fig. 3I). Consistent with the murine data, patients with TR exhibited increased plasma IL-6 (Fig. 3J). Further, flow cytometry confirmed a higher abundance of IL-6+ CD14+ CD16+ monocytes compared to controls (Fig. 3K). Systemic IL-6 is a well-established therapeutic target in cardiovascular disease (*25, 26*); however, its association with congestion has not been clearly defined. To address this, we measured systemic IL-6 levels across different murine chronic HF models representing both forward and backward failure. Notably, IL-6 was predominantly upregulated in the TR model (fig. S7F). These findings were corroborated in a human cohort (*27*), where LV ejection fraction showed only a modest correlation with circulating IL-6 levels. In contrast, the presence of edema - used as a surrogate marker of congestion - emerged as a significantly stronger predictor of elevated IL-6 (fig. S7G–H). Together, these data indicate that chronic venous congestion in TR drives systemic IL-6 upregulation, with peripheral monocytes serving as a major source.

### Gut-derived endotoxemia reprograms circulating monocytes to drive TR-induced cardiac fibrosis

To test whether TR-induced myeloid alterations causally contribute to adverse cardiac remodeling, we isolated circulating and splenic monocytes from mice one month after TR or sham intervention and adoptively transferred them into recipient mice with prior IR injury, modeling pre-existing left-sided HF (Fig. 4A, fig. S8A–B). Monocyte transfer was initiated 10 days after IR injury and repeated every other day for four weeks to ensure sustained exposure. Recipient mice were stratified by infarct size before group assignment (Fig. 4B). At study endpoint, recipient mice with pre-existing left-sided HF that received monocytes from TR donors exhibited significantly increased heart-to-body weight ratios, indicative of aggravated chronic cardiac remodeling (Fig. 4C). Echocardiographic assessment further demonstrated a significant deterioration in LV systolic function in mice receiving TR-derived monocytes (Fig. 4D–E). Histological analysis of recipient hearts uncovered significantly increased left ventricular fibrosis following transfer of monocytes from TR donors compared to sham donors (Fig. 4F–G). Collectively, these results propose that monocytes are not only activated and reprogrammed in TR, but that these changes in the myeloid compartment are sufficient to accelerate left-sided HF.

**Fig. 4.**
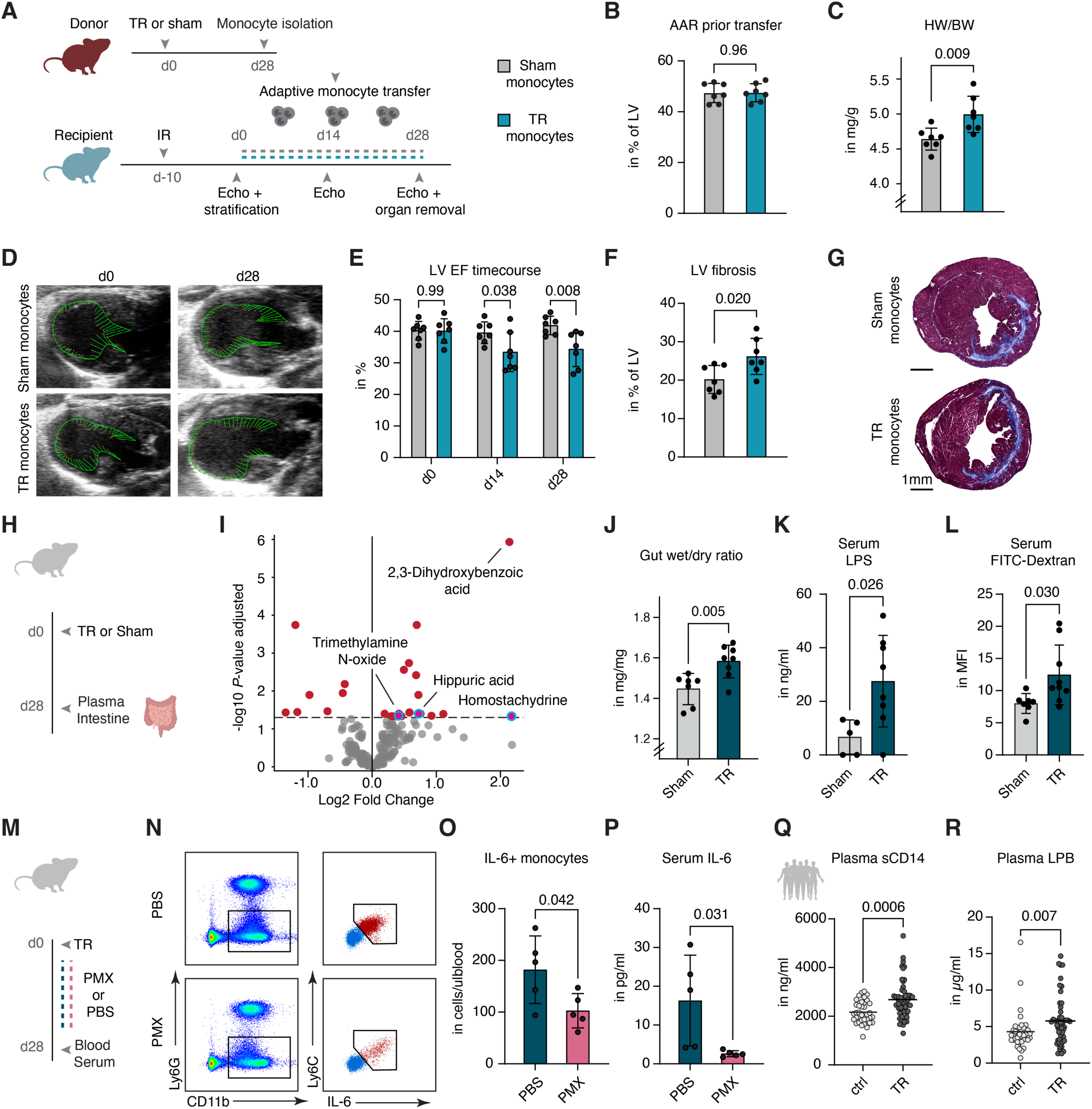
TR-induced cardiac remodeling is monocyte-dependent and driven by gut-derived endotoxemia. (**A**) Monocytes isolated from TR or sham donor mice (red) at d28 were adoptively transferred into recipient mice (blue) with IR-induced left-sided HF every other day from d0 through d28. (**B**) Pre-stratification of recipient mice by echocardiographic assessment of area-at-risk (AAR) prior to monocyte transfer (n = 7, unpaired t-test, mean ± SD). (**C**) Heart-to-body weight ratio (HW/BW) at d28 (n = 7, unpaired t-test, mean ± SD). (**D**) Representative echocardiographic B-mode images of recipient LV at indicated timepoints; green lines denote endocardial movement. (**E**) Echocardiographic evaluation of LV EF in mice during monocyte transfer experiments (n = 7, two-way ANOVA with Sidak’s multiple comparison, mean ± SD). (**F**) Quantification of LV fibrosis from short-axis histological sections at d28 (n = 7, unpaired t-test, mean ± SD). (**G**) Representative Masson’s trichrome-stained short-axis sections at d28. Scale bar, 1mm. (**H**) Schematic of the experimental setup. Blood and intestinal tissue were collected at d28 from TR and sham mice for plasma metabolomics, LPS quantification, and intestinal assessment. (**I**) Volcano plot of plasma metabolites in TR vs. sham mice on d28; red points indicate *P*adj. < 0.05 (n = 8, Welch’s t-test, Benjamini-Hochberg correction). (**J**) Intestinal wet-to-dry weight ratio as a measure of tissue edema (n = 7-8, unpaired t-test, mean ± SD). (**K**) Circulating LPS levels in serum at d28 (n = 5-7, unpaired t-test, mean ± SD). (**L**) Gut permeability assessed by FITC-dextran assay following oral gavage and 4-hour serum collection (n = 6-9, unpaired t-test, mean ± SD). (**M**) Schematic depicting intraperitoneal administration of Polymyxin B (PMX) or PBS (control) following TR induction, with blood collection at d28 for flow cytometry and serum IL-6 measurement. (**N**) Representative flow cytometry plots showing IL-6+ monocytes in TR+control (top) vs. TR+Polymyxin B (PMX; bottom). (**O**) Frequency of IL-6+ monocytes based on flow cytometry in TR mice at d28 treated with control or PMX (n = 5, unpaired t-test, mean ± SD). (**P**) Serum IL-6 levels in TR mice at d28 treated with control or PMX (n = 5, unpaired t-test, mean ± SD). (**Q-R**) Plasma levels of soluble CD14 (Q) and LPS-binding protein (LBP) (R) in TR patients vs. controls (n = 50 TR, n = 35 controls, Mann-Whitney test, mean ± SD).

We then sought to identify the upstream drivers of monocyte reprogramming in TR. Building on the enriched gene programs in monocytes in our human scRNA-seq analysis (fig. S2J–K), we explored the splanchnic compartment as a potential trigger of myeloid activation. Unbiased plasma metabolomic profiling in TR mice revealed enrichment of gut-derived metabolites (Fig. 4H-I), corroborating this hypothesis. Inspection of the intestine revealed macroscopic abnormalities (fig. S8C-D) and increased tissue edema, as reflected by a higher wet-to-dry ratio (Fig. 4J). Immunofluorescence staining further revealed a marked reduction in the tight junction protein ZO-1 in intestinal tissue of TR mice compared to sham controls (fig. S8E–F), indicative of compromised intestinal barrier integrity. Consistent with increased intestinal permeability, mice with TR exhibited elevated levels of circulating LPS (Fig. 4K). This was further supported by a FITC-dextran assay, in which oral administration of fluorescent dextran resulted in increased plasma fluorescence in TR mice, indicating enhanced luminal-to-vascular translocation (Fig. 4L). Pharmacological neutralization of LPS with Polymyxin B markedly reduced the proportion of IL-6+ monocytes and decreased systemic IL-6 levels (Fig. 4M–P), identifying gut-derived endotoxemia as a key upstream trigger of monocyte activation in TR. In line with these findings, patients with TR displayed increased circulating levels of LPS-binding protein (LBP) and soluble CD14 (Fig. 4Q–R), both established markers of microbial translocation (*28*). Together, these findings demonstrate that TR-induced venous congestion disrupts intestinal barrier integrity, triggering gut-derived endotoxemia that reprograms circulating monocytes and drives secondary cardiac fibrosis independent of direct hemodynamic stress.

### IL-6 blockade rescues TR-associated biventricular remodeling

Our results raise the possibility of utilizing IL-6 blockade therapeutically to prevent TR-mediated HF progression. To explore this, we performed antibody-mediated IL-6 blockade in mice subjected to combined IR injury and TR (Fig. 5A). Consistent with our prior experimental design, TR was induced 10 days after MI to avoid interference with the acute inflammatory phase, during which IL-6 is known to play a critical role (*29*). Animals were stratified according to infarct scar size and TR severity prior to treatment (Fig. 5B–C). Remarkably, IL-6 inhibition significantly improved LV ejection fraction and attenuated TR-induced adverse remodeling (Fig. 5D–E). Histological analyses demonstrated a marked reduction in left ventricular fibrosis (Fig. 5F–I). At the cellular level, snRNA-seq demonstrated a significant decrease in fibroblast abundance (Fig. 5J–K). These compositional alterations were paralleled by broad transcriptomic normalization across fibroblasts and myeloid cells (Fig. 5L–M), including downregulation of pro-fibrotic programs (e.g. Postn, Ccn1, Ccn4, Ccn5, Runx1). Notably, IL-6 inhibition also improved right ventricular performance and right ventricular fibrosis in the setting of pre-existing left-sided HF with superimposed TR (Fig. 5N–P). Collectively, these data reveal the therapeutic potential of IL-6 inhibition in chronic TR.

**Fig. 5.**
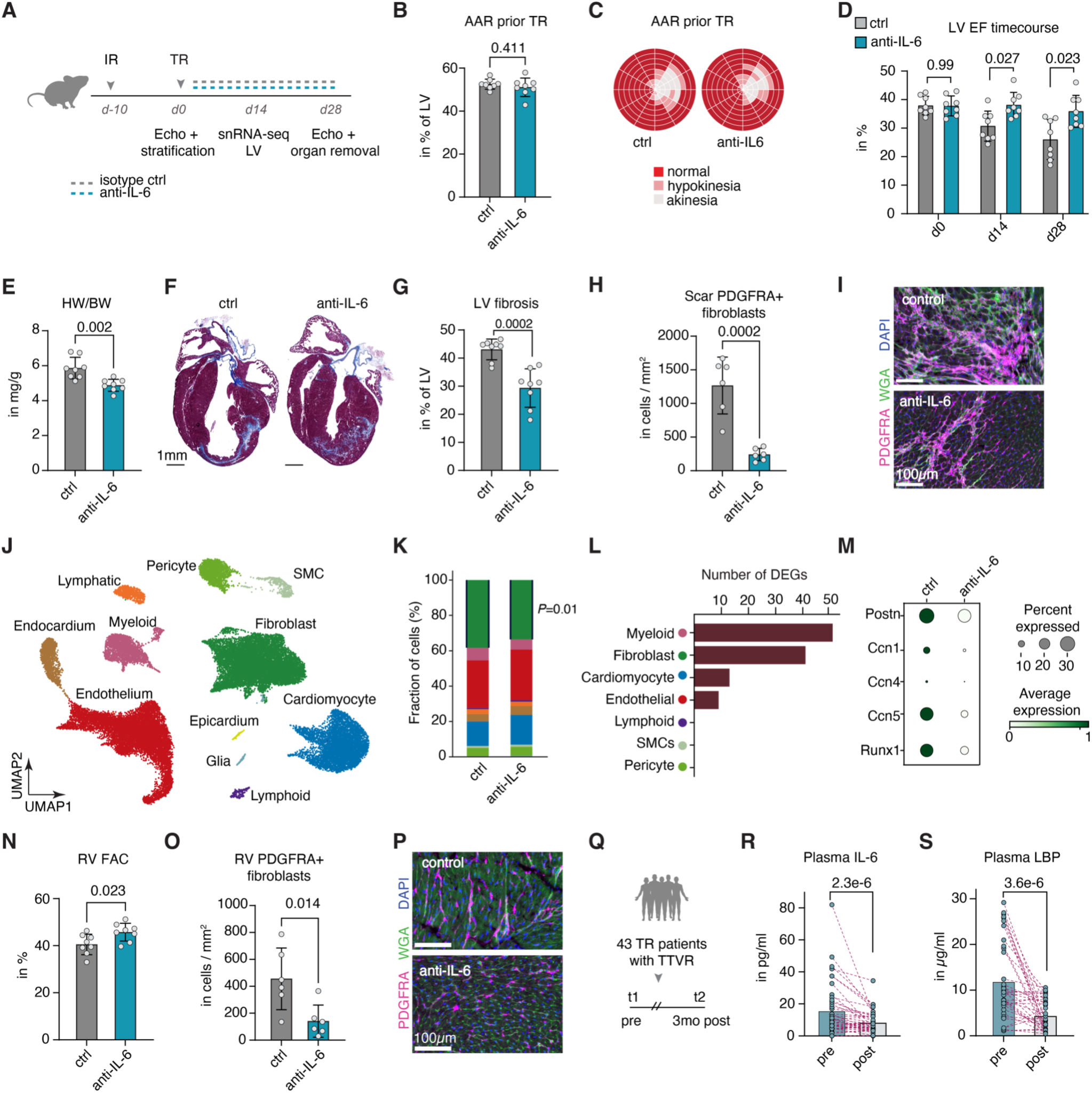
IL-6 blockade improves TR-associated biventricular remodeling and cardiac dysfunction. (**A**) Schematic experimental design. Mice underwent IR injury at day −10, followed by TR induction at day 0. Anti-IL-6 antibody or control treatment was administered continuously from day 0 through day 28, with assessments on day 14 and 28. (**B**) Area-at-risk (AAR) prior to TR induction in anti-IL-6 and control groups (n = 8, unpaired t-test, mean ± SD). (**C**) Representative bull’s eye plots depicting akinetic/hypokinetic LV at baseline stratification. (**D**) LV EF in control and anti-IL-6 mice at days 0, 14, and 28 post-TR (n = 8, two-way ANOVA with Sidak’s multiple comparison, mean ± SD). (**E**) Heart-to-body weight ratio at day 28 (n = 8, unpaired t-test, mean ± SD). (**F**) Representative Masson’s trichrome sections from control (left) and anti-IL-6 (right) hearts at day 28. Scale bar, 1mm. (**G**) LV fibrosis at d28 quantified as percentage of area-at-risk (n = 8, unpaired t-test, mean ± SD). (**H**) PDGFRA+ fibroblast density in the LV scar (n = 6, unpaired t-test, mean ± SD). (**I**) Immunofluorescence stainings of cardiac cross-sections showing PDGFRA (magenta), WGA (Wheat Germ Agglutinin, green), and DAPI (blue) in control (top) and anti-IL-6 (bottom) hearts 4 weeks post-TR. Scale bar, 100µm. (**J**) UMAP of LV nuclei from mice with IR+TR, with or without anti-IL-6 treatment, profiled by snRNA-seq at day 14 post-TR. Colors denote annotated cell types. SMC, smooth muscle cell. (**K**) Relative cell-type proportions per condition (WLS regression, Benjamini-Hochberg correction, *P*-value shown for Fibroblast). (**L**) Number of DEGs per cell type between conditions, determined by pseudobulk DESeq2 (n = 3 per group, Wald test, Benjamini-Hochberg, *P*adj. < 0.05). (**M**) Dot plot depicting the expression of indicated genes in fibroblasts based on snRNA-seq analysis. (**N**) RV fractional area change (FAC) at day 28 (n = 8, unpaired t-test, mean ± SD). (**O**) PDGFRA+ fibroblast density in RV tissue at d28 post-TR (n = 6, unpaired t-test, mean ± SD). (**P**) Representative immunofluorescence stainings of RV cross-sections 4 weeks after IL-6 blockade or control treatment; PDGFRA (magenta), WGA (green), DAPI (blue). (**Q**) Study design for human validation. 43 patients from two centers undergoing TTVR (Transcatheter Tricuspid Valve Repair, including edge-to-edge repair or valve replacement) with significant TR reduction were included. **(R)** Serial plasma IL-6 measurements before and after TTVR (Wilcoxon matched-pairs signed rank test, mean with individual data points, lines connect paired samples). **(S)** Plasma LBP levels before and after TTVR (Wilcoxon matched-pairs signed rank test, mean with individual data points, lines connect paired samples).

### Transcatheter-repair impacts IL-6 levels in patients with TR

To assess whether TR-induced IL-6 elevation is reversible upon hemodynamic correction, we analyzed serum from 43 individuals undergoing transcatheter tricuspid valve repair (TTVR), including edge-to-edge repair or valve replacement, restricting the analysis to procedural responders (Fig. 5Q). Successful intervention led to a significant reduction in circulating IL-6 levels (Fig. 5R, table S4), paralleled by decreased plasma levels of LPS-binding protein (LBP) (Fig. 5S). In summary, these data establish IL-6 as a central mediator linking TR-induced myeloid activation to adverse biventricular remodeling and identify the IL-6 axis as a modifiable driver of HF progression in TR.

## DISCUSSION

Right HF represents a major unmet clinical challenge, driven by the absence of targeted therapies and an incomplete understanding of its molecular underpinnings. Here, we present a cross-species analysis integrating patients and mouse models of left and right ventricular failure with TR, uncovering mechanistic insights with direct translational potential. Using single-cell and single-nucleus transcriptomic approaches alongside functional in vivo experiments, we demonstrate that TR drives marked expansion and inflammatory reprogramming of circulating monocytes, which in turn worsen pre-existing dysfunction of the LV.

The causal role of right ventricular dysfunction and secondary TR in left-sided heart disease has been long debated, with questions of whether TR is merely a bystander or an active contributor (*30*). Our work provides mechanistic evidence that superimposed TR accelerates left ventricular remodeling and functional decline. Although hemodynamic factors likely contribute to this process, adoptive transfer experiments position inflammation as the central driver of the maladaptive crosstalk between the failing right and left ventricles. Importantly, IL-6 emerges as a key effector in this process. Although elevated IL-6 levels have been widely reported in HF (*21, 27*), our data identify the congestive component associated with TR as a critical trigger for IL-6 induction. This finding refines the current understanding of inflammatory signaling in HF by implicating venous congestion and right-sided valvular dysfunction as one upstream driver of IL-6-mediated pathways. Consistent with this mechanism, therapeutic blockade of IL-6 in our preclinical model attenuated cardiac fibrosis and improved functional outcomes, supporting the concept that targeting IL-6 signaling may represent a promising therapeutic strategy in this context. Notably, the IL-6 ligand inhibitor Ziltivekimab is currently under clinical investigation in HF through the ongoing HERMES (NCT05636176) and ATHENA (NCT06200207) trials. Our data suggest that patients with concomitant venous congestion and right-sided valvular dysfunction may represent a particularly responsive subgroup, in whom gut-derived endotoxemia and IL-6 induction may serve as mechanistic drivers of disease progression.

Collectively, these findings integrate hemodynamic, cellular, and molecular insights to redefine TR as more than a passive consequence of advanced disease, instead identifying it as an active driver of immune-mediated cardiac remodeling. By linking valvular hemodynamics to systemic immune activation and ventricular fibrosis, our work uncovers a mechanistic axis through which right heart dysfunction accelerates left ventricular failure. These results establish a conceptual framework in which targeting both structural abnormalities and inflammation may be required to interrupt the vicious cycle of biventricular deterioration in HF. Ultimately, this study highlights immunomodulation - alongside hemodynamic and structural interventions - as a promising strategy to prevent or even reverse the progression of complex right and left ventricular HF.

## MATERIALS AND METHODS

### Ethical approval for human specimens

The study was conducted in accordance with the Declaration of Helsinki and was approved by the University Hospital Heidelberg Review Board (approval nos. S-738/2023, S-277/2024, S-018/2022) and University Medical Center Schleswig-Holstein Review Board (approval no. D506/22 - AZ F02/24). Written informed consent was obtained from all patients prior to enrollment. All data were de-identified prior to analysis. Demographic and clinical characteristics of all patients are provided in table S1-S4.

### Retrospective observational cohort study

Patients were included if they had available baseline and follow-up echocardiographic data and a baseline LV EF < 50%. The TR group was defined as patients with TR severity grade ≥ 3 at both baseline and follow-up (persistent severe/torrential TR), while the control group comprised patients with TR severity grade 1 at baseline and non-severe TR at follow-up. Patients meeting neither criterion were excluded from the analysis. To reduce confounding, propensity scores were estimated for each patient using multivariable logistic regression with age, sex, and baseline LV EF as covariates. Patients in the TR and control groups were then matched in a 1:4 ratio (TR:control) using nearest-neighbor propensity score matching without replacement. Covariate balance before and after matching was assessed using standardized mean differences (SMD), with SMD < 0.1 considered indicative of adequate balance. In the matched cohort, follow-up LV EF was analyzed using multivariable linear regression and all-cause mortality was analyzed using multivariable logistic regression, both adjusting for TR group assignment, age, sex, and baseline LV EF. All analyses were performed in Python using the statsmodels and scikit-learn libraries.

### Human plasma protein profiling

Blood was collected into EDTA-containing tubes and centrifuged at 2000 x g for 10 minutes at room temperature. The plasma fraction was transferred to a new tube and stored at −80 °C until analysis. Plasma protein profiling was conducted using the Olink Target 96 Inflammation panel (Olink Proteomics) comprising 92 protein biomarkers. Sample processing and measurements were conducted by the Microarray Core Facility at the German Cancer Research Center (DKFZ, Heidelberg, Germany) in accordance with the manufacturer’s instructions. Plasma protein abundances were measured using the proximity extension assay (PEA) principle, in which pairs of oligonucleotide-labeled antibodies bind target proteins and, upon proximity, generate amplifiable DNA reporters quantified by qPCR. Manufacturer-provided internal controls were included in each assay run to monitor assay performance and detection efficiency. Protein expression levels were reported as Normalized Protein Expression (NPX) values, representing relative, log2-transformed units. Quality control and normalization were performed using the NPX Signature Software V2.0.2. Downstream data analysis was carried out using the Olink Stat Analyzer software. Proteins with an adjusted *P*-value of <0.05 were considered to be differentially expressed. Plasma IL-6 concentrations in participants from the TTVR cohort were quantified using the Milliplex Human Cytokine/Chemokine Magnetic Bead panel (Luminex, cat. no. HCYTA-60K-16), the Olink Target 48 Cytokine panel (Olink Proteomics), and the Human IL-6 Quantikine ELISA Kit (R&D Systems, cat. no. D6050B). For the Olink panel, protein concentrations were calculated using manufacturer-provided calibrators. All three platforms yielded concordant absolute concentrations (pg/ml). Plasma concentrations of sCD14 and LBP were quantified by ELISA using the Human CD14 Quantikine ELISA Kit (R&D Systems, cat. no. DC140) and the Human LBP ELISA Kit (Abcam, cat. no. ab213805), respectively, according to the manufacturer’s instructions.

### Human PBMC isolation and fixation

For each sample, 4 mL of freshly collected whole blood in EDTA tubes were mixed with 4 mL of 1x RoboSep Buffer (STEMCELL Technologies, #20124) and carefully layered over a Lymphoprep density gradient (STEMCELL Technologies, # 18061). Samples were centrifuged at 1200 x g for 10 minutes with the brake enabled. PBMCs were then washed twice, and the resulting pellet was resuspended in 2 mL of RoboSep Buffer. PBMCs were counted in a Neubauer hemocytometer, and 1 million cells were transferred to Evercode Cell Fixation v3 kit (Parse Biosciences, ECFC3300) according to the Parse Biosciences Cell Fixation protocol. In brief, cells were fixed and permeabilized using the proprietary fixation and permeabilization reagents, followed by quenching of the reactions and resuspension in a cryoprotective storage buffer containing DMSO. Fixed cells were filtered, counted, and cryopreserved at −80 °C using controlled-rate freezing prior to downstream Evercode library preparation.

### Human scRNA-seq library preparation

Collected fixed PBMCs were processed using the Evercode WT Mega v3 kit (Parse Bioscience, ECWT3500). The libraries were converted for sequencing on the Ultima UG100 instrument. After sequencing a raw read trimming step (analysis: scRNA_Parse_v3) was applied to the base call files to generate paired end fastqs files (R1: 90bp, R2: 58bp) for Trailmaker analysis. During this step, reads lacking valid Parse barcodes were filtered out at the sequencer level.

### Human scRNA-seq analysis

FASTQ files were processed, aligned to the reference genome (GRCh38), and quantified using the Parse Trailmaker pipeline following the manufacturer’s recommendations. Raw count matrices, gene annotations, and cell metadata were assembled into AnnData objects using Scanpy (v1.12.1). Genes lacking valid gene names were removed prior to analysis. Cells expressing fewer than 200 genes and genes detected in fewer than 3 cells were excluded. Cells with >5% mitochondrial counts were removed. Doublets were detected per sample using Scrublet (expected rate 0.05; score threshold >0.2) and excluded. Counts were normalized, log-transformed, and highly variable genes were selected per sample. After PCA, batch correction was applied using Harmony. A k-nearest-neighbor graph (k=16) was used for UMAP embedding and Leiden clustering (resolution 0.75). Clusters were manually annotated to 13 cell types. Cell-type composition differences between TR and controls were tested by weighted least squares regression with Benjamini-Hochberg correction (*31*). Differential gene expression was assessed by Wilcoxon rank-sum test, followed by pre-ranked gene set enrichment analysis against Gene Ontology (GO) Biological Process 2025. For the validation cohort, a pre-processed gene expression matrix from a published PBMC scRNA-seq dataset was used (*16*). The dataset was first subset to include only patients with HFrEF. Clinical metadata from the source study were subsequently integrated with patient-level records to identify the presence and echocardiographic severity of TR. Patients were stratified into three groups: no TR, mild TR, and moderate-to-severe TR. Reanalysis was performed in R using the Seurat package. Cell types were retained from the original publication, where annotation had been based on canonical marker gene expression, identifying monocytes, T cells, NK cells, progenitor cells, B cells, and megakaryocytes. The relative proportions of each cell population were calculated per patient and visualized as stacked bar charts grouped by TR severity

### Analysis of human snRNA-seq data

Processed snRNA-seq data (*14*) were reanalyzed after incorporating additional clinical information on the presence of clinically relevant TR using R and the R package Seurat. Cell type annotations were derived from the original study and were validated using canonical gene marker genes. Cell-type composition differences between TR and non-TR samples were quantified per donor as the percentage of total captured nuclei, and statistical differences were assessed using weighted least squares regression with TR status as the predictor. Differentially expressed genes between conditions were calculated using the Wilcoxon rank-sum test via FindMarkers function. Gene set enrichment analysis was performed using Enrichr via the gseapy Python package, testing the top genes for enrichment in GO Biological Process 2023 terms. Enriched pathways were identified with a significance cutoff of 0.05. Fibroblasts were extracted and re-clustered independently. The SCTransform-normalized count matrix was used to compute PCA (50 components), a shared nearest-neighbor graph was constructed on the first 20 principal components, and Leiden-equivalent graph-based clustering was applied at resolution 0.3, followed by UMAP for visualization. Low-quality fibroblast subclusters were removed, and the remaining cells were annotated into five subtypes (Fib1–Fib5) based on subcluster-specific marker genes.

### Mice

C57BL/6N mice were obtained from Janvier Labs (Saint-Berthevin, France) and were studied at 10 to 20 weeks of age unless otherwise specified. Animals were housed under standard laboratory conditions with a 12-hour light/12-hour dark cycle and ad libitum access to water and food. All experiments with isolated TR induction were performed in male and female mice at equal sex ratios. No sex-dependent differences were observed in the primary outcomes assessed. For combined IR and TR experiments, including those incorporating snRNA-seq analyses, female mice were used exclusively to minimize biological variability. All animal procedures were approved by the institutional review board of the University of Heidelberg, Germany, and the responsible government authority of Baden-Württemberg, Germany (project number G-94/21 and G-23/22).

### Murine TR induction

30 minutes prior to procedure start, mice received a single dose of subcutaneous buprenorphine (0.05 mg/kg) for analgesia. Real-time imaging during the procedure was performed using a Vevo 2100, Vevo 3100, or Vevo F2 system equipped with an MS550D, MX400 or MX550D transducer, respectively (Fujifilm VisualSonics, Toronto, Canada). For preparation, the fur on the left chest was removed with a chemical hair remover. Mice were anesthetized with 1.5-2% isoflurane and positioned supine on the imaging platform. Cardiac function was briefly evaluated, followed by visualization of the tricuspid valve in B-mode and color Doppler mode in a modified parasternal short-axis imaging. A 27-gauge microscissors (BVI Medical, SC27.D03), mounted on an additional micromanipulator, was aligned parallel to the transducer. The microscissor, closed using a custom clip, were inserted into the left lateral chest under image guidance and advanced to the tricuspid valve via transapical, transseptal puncture. Correct placement was confirmed by the presence of a regurgitant jet in color Doppler mode. The microscissors were then opened, and the tricuspid valve was severed repeatedly until a robust regurgitant jet was observed. Upon completion, the microscissors were retracted through the left ventricular myocardium under continuous image guidance. The closed tip was maintained in a mid-myocardial position during withdrawal for 2-3 seconds to allow endocardial sealing and prevent pericardial effusion. Successful TR induction was confirmed by immediate RA and RV dilation and characteristic regurgitant jet on color Doppler. For sham procedure, the microscissors were advanced into the RV via the same transapical, transseptal approach, but remained closed and were retracted without valve severing. Absence of interventricular shunting was confirmed in each animal by color Doppler imaging.

### Echocardiography-guided IR induction

Echocardiography-guided ischemia/reperfusion (EG-IR) was performed as described previously (*17*). In brief, mice received subcutaneous buprenorphine (0.05 mg/kg) for analgesia 30 minutes prior to procedure onset. Mice were anesthetized with 1.5% isoflurane and positioned supine on the imaging platform. The heart was visualized in a modified short axis view. Coronary arteries were scanned from base to midpapillary level using color Doppler mode. Two 30-gauge needles, each mounted on additional micromanipulators, were inserted percutaneously and positioned ventral and dorsal to the LAD. The needle tips were advanced toward one another to compress the LAD. Successful occlusion was confirmed by loss of Doppler flow signal, regional myocardial akinesia, and ST elevation in the ECG. Occlusion was maintained for 45 minutes to generate mid-size infarcts. Five minutes prior to reperfusion, long-axis imaging was acquired to quantify the area-at-risk to determine akinetic ischemic area. Reperfusion was achieved by separating and removing the needles. Restoration of coronary flow was confirmed by return of Doppler signal and post-ischemic changes, such as wall thickening and myocardial edema.

### Echocardiography

Transthoracic echocardiography was performed using a Vevo 2100, Vevo 3100, or Vevo F2 high-frequency ultrasound system equipped with MS550D, MX400 or MX550D transducers, respectively (Fujifilm VisualSonics, Canada). Mice were anesthetized with 1.0% to 1.5% isoflurane in oxygen and placed on a temperature-controlled platform. Standard parasternal long axis views of the LV and the RV were obtained, followed by parasternal short-axis views at the mid-papillary level. Modified short-axis views were acquired for visualization of the RV and the RA. IVC imaging was additionally performed in a subcostal long-axis and short axis view to assess venous congestion. Echocardiographic measurements were performed in a blinded manner if applicable using VevoLab software (Fujifilm VisualSonics, Canada). Measurements of wall thickness were performed at end-diastole. For noninvasive measurement of the area at risk (AAR), the akinetic fraction of the LV was determined as the percentage of akinetic traced endocardial length relative to the total traced endocardial length.

### Monocyte depletion and monocyte transfer

To deplete CCR2+ monocytes, mice were given 10 μg anti-CCR2 rat monoclonal antibody MC21 or control antibody (BioXCell, clone LTF-2) i.p. for five consecutive days. For monocyte transfer experiments, mice were euthanized, spleens were harvested and stored in phosphate-buffered saline (PBS), and blood was collected into labeled EDTA-coated tubes that were gently inverted to prevent clot formation. Spleens were mechanically dissociated by passage through a 40 µm cell strainer using a syringe plunger in 20 mL of 1× RoboSep buffer (Stemcell Technologies), followed by centrifugation at 400 × g for 5 minutes at 4 °C. Whole blood was diluted tenfold in 1× red blood cell (RBC) lysis solution and incubated for 20 minutes. Single cell suspensions of spleens were incubated in 2 ml RBC lysis solution for 2 minutes. After completion of lysis, samples were quenched with 20 mL RoboSep buffer, centrifuged at 400 × g for 5 minutes at 4 °C, resuspended in 1 mL RoboSep buffer, and processed for monocyte isolation using the EasySep Mouse Monocyte Isolation Kit according to the manufacturer’s instructions. Isolated monocytes were resuspended in RPMI-1640 medium (Sigma-Aldrich), and 2 × 10⁵ cells in 100 µL were administered per mouse via tail vein injection.

### In vivo IL-6 blockade and LPS blockade

To neutralize IL-6 signaling in vivo, mice received intraperitoneal injections of anti-mouse IL-6 antibody (InVivoMAb clone MP5-20F3, BioXCell, cat. #BE0046) at 400 µg per dose, three times weekly for 2 weeks (snRNA-seq analysis) or 4 weeks (functional assessment). Rat IgG1 isotype control antibody (anti-horseradish peroxidase, BioXCell, cat. #BE0088) was administered at the same dose and schedule. For LPS neutralization, mice received intraperitoneal injections of polymyxin B at 50 µg per 100 µl sterile PBS three times weekly for 4 weeks. Vehicle-treated control animals received 100 µl sterile PBS.

### Histopathological analysis

Hearts were rinsed in PBS and sectioned along the long or short axis using a scalpel. Tissues were embedded in Tissue-Tek O.C.T. Compound (Sakura) and frozen in 2-methylbutane (Honeywell) on dry ice. Organs were cryosectioned at 8-9 µm on a cryostat (Leica Biosystems, Germany), collected on adhesion slides (Epredia SuperFrost Plus, Thermo Fisher Scientific) and air-dried at room temperature for 1h. Sections were stored at −80°C until use. Cardiac or hepatic fibrosis was evaluated using Picrosirius Red staining (Abcam, ab245887) or Masson’s Trichrome staining (Sigma-Aldrich, HT15-1KT) according to the manufacturer’s protocols. Stained sections were mounted and imaged using a slide scanner (Zeiss, Axioscan 7). Whole-section images were manually annotated into the following anatomical regions of interest: area at risk (defined as the transmural region exhibiting mid-myocardial scarring), LV free wall, LV septal wall, RV, right atrium, and left atrium. For fibrosis quantification, a pixel classifier was trained on representative images spanning each experimental series and applied to classify fibrotic and non-fibrotic areas based on Masson’s Trichrome staining intensity. Fibrosis was expressed as the percentage of fibrotic area relative to the total annotated region area.

### ELISA measurements

For serum measurements, peripheral blood was collected from facial vein puncture in a MiniCollect CAT Serum Separator tube (Greiner Bio-One, Austria, cat. no. 450533) at indicated time points. Serum was obtained after centrifugation (4°C, 2000g, 10 minutes). For Troponin T measurements after EG-IR, serum was 40 times diluted in PBS and cTnT was detected using an automated Cobas Troponin T hs STAT Elecsys (Roche). Serum levels of IL-6 were analyzed using a Mouse IL-6 Quantikine ELISA Kit (R&D Systems, M6000B) following the manufacturer’s instructions. Serum levels of mouse LPS were analyzed using Mouse LPS ELISA Kit (MyBioSource MBS261904).

### Mouse serum Olink cytokine assay

Serum was collected via facial vein puncture, and cytokines were measured using the Olink Target 48 Mouse Cytokine panel (Olink Proteomics) following the manufacturer’s protocol. 3-4 biological replicates were analyzed per group. Protein abundances were reported as Normalized Protein Expression (NPX) values on a log2 scale, following internal normalization. Only assays that passed Olink quality control were included in the downstream analyses. Statistical significance was assessed using Welch’s t-test, followed by Benjamini-Hochberg correction for multiple testing. For validation of cytokines in a separate cohort, cytokine concentrations were quantified using the Milliplex MCYTOMAG-70K-16C magnetic bead-based multiplex assay (Millipore, Sigma-Aldrich) according to the manufacturer’s instructions. Briefly, samples were processed in duplicate, incubated with antibody-coated magnetic beads, and detected using streptavidin-phycoerythrin. Data were acquired on a Luminex MAGPIX system and analyzed using xPONENT software. Cytokine concentrations were calculated from standard curves and samples with concentrations below the lower limit of detection were assigned the minimum detectable value.

### Murine nuclei isolation and snRNA-seq library preparation

Snap-frozen cardiac tissue samples were transferred into 2 ml of ice-cold Nuclei Extraction Buffer (Miltenyi Biotec, #130-128-024) buffer supplemented with 0.2 U/µl RNAse inhibitor (Invitrogen, AM2696) in gentleMACS C Tubes (Miltenyi Biotec, #130-093-237). Tubes were placed on a gentleMACS Octo Dissociator (Miltenyi Biotec, #130-095-937), and the 4C_nuclei_1 program was run. After program termination, the nuclei suspension was applied to 30 µm MACS SmartStrainers (Miltenyi Biotec, #130-098-458) and washed with 2 ml ice-cold Nuclei Extraction Buffer. Samples were centrifuged at 300 x g at 4°C for 10 minutes. The nuclei pellet was resuspended in 450 µl Nuclei Separation Buffer (0.04% final BSA, 14% Nuclei Extraction in PBS) and 50 µl of Anti-Nucleus MicroBeads (Miltenyi Biotec, #130-132-997) were added and incubated for 15 minutes at 4°C. Two milliliters of Nuclei Separation Buffer were added, and the suspension was applied to pre-cooled and pre-rinsed LS columns (Miltenyi Biotec, #130-042-401) placed on a QuadroMACS Separator. Columns were washed twice with 1 ml of Nuclei Separation Buffer, removed from the separator, and placed on a collection tube to elute magnetically labeled nuclei with 1 ml of Nuclei Separation Buffer. Nuclei were counted using propidium iodide staining on a fluorescence cell counter (Logos Biosystems, Luna FL). One million nuclei were transferred to Evercode Nuclei Fixation v3 (Parse Biosciences, ECFN3300) and processed according to the manufacturer’s protocol. Fixed nuclei were cryopreserved at −80 °C until library preparation. Single-nuclei libraries were prepared using the Parse Evercode Whole Transcriptome Kit v3 according to the manufacturer’s instructions and sequenced on an Illumina NextSeq 2000.

### Murine snRNA-seq analysis

FASTQ files were processed, aligned to the reference genome (GRCm39), and quantified using the Parse Trailmaker pipeline following the manufacturer’s recommendations. Raw count matrices were assembled into an AnnData object using Scanpy (v1.12.1). Cells with fewer than 200 detected genes, genes present in fewer than 3 cells, and cells with >5% mitochondrial reads were excluded. Doublets were identified and removed per sample using Scrublet (expected doublet rate = 0.05). Counts were library-size normalized and log1p-transformed. Highly variable genes were selected in a batch-aware manner, and PCA was performed on scaled data. Sample-level batch effects were corrected using Harmony. A k-nearest-neighbor graph (k = 30) was constructed on the corrected embedding and visualized by UMAP. Leiden clustering (resolution = 0.75) was applied and cluster marker genes identified by Wilcoxon rank-sum test. Clusters were manually annotated into 11 cardiac cell types based on canonical marker expression. Clusters lacking clear marker gene identity were excluded as contamination. Cell type proportions were calculated per sample and compared between conditions using weighted least squares regression, with Bonferroni correction for multiple comparisons. Fibroblasts were subset and reprocessed independently from raw counts, followed by Leiden clustering (resolution = 0.3) and UMAP visualization. Pseudobulk differential expression was performed per cell type using DESeq2 (pydeseq2), comparing IR+TR to IR+sham. Genes were pre-filtered by minimum expression and sample-level proportion thresholds using decoupler. Wald tests were applied and results reported as log2 fold changes with adjusted *P*-values. Gene set enrichment analysis was performed using GSEApy prerank with the GO Biological Process 2025 gene set library. Genes were ranked by the product of −log10(adjusted *P*-value) and log2 fold change.

### Statistics and reproducibility

For treatment cohorts and combined IR/TR studies, group assignment was performed using an iterative randomization procedure to minimize variance in group means of infarct scar size and vena contracta width. For all other experiments, mice were randomly assigned to experimental groups. Animals with mild TR in whom no corresponding increase in IVC diameter was detected were excluded from further experiments. Sample size calculation was performed in G*Power 3.1 in accordance with the approved animal protocol and was based on our experience with similar experimental studies to achieve 80% power at a significance level of P < 0.05. All analyses were performed blinded to experimental group allocation. Statistical analysis and graphical representation were performed using RStudio and GraphPad Prism software (version 10; GraphPad, San Diego, CA). Shapiro-Wilk test was used to assess normal distribution of the data. Normally distributed data were analyzed by Student’s two-tailed unpaired or paired t-test, one-way or two-way ANOVA with multiple comparison correction by Tukey’s test or Sidak’s test, as appropriate. Non-normally distributed data were compared using Mann-Whitney test for two independent groups, Wilcoxon matched-pairs signed rank test for two paired groups, and Kruskal-Wallis test for more than two groups. *P* < 0.05 was considered statistically significant. Data are expressed as mean ± SD with individual data points as indicated, respectively.

## List of Supplementary Materials

Materials and Methods

Figs. S1 to S8

Tables S1 to S4

References (*32*)

Movie S1

