## Supplementary for "Tricuspid valve regurgitation accelerates heart failure via a cardio-intestinal innate immune circuit"

**The PDF file includes:**

Materials and Methods
Figs. S1 to S8
Tables S1 to S4
References

**Other Supplementary Materials for this manuscript include the following:**

Movie S1

### Materials and Methods

Analysis of the BIOSTAT-CHF Human Cohort

To examine the association between IL-6 and right-sided congestion, we leveraged data from the BIOSTAT-CHF index cohort (*32*). Briefly, BIOSTAT-CHF was a prospective, multi-center cohort enrolling patients from 11 European countries. Participants were aged ≥18 years and presented with new-onset or worsening symptoms of HF, combined with an LVEF ≤40% and/or BNP >400 pg/mL or NT-proBNP >2000 pg/mL. Plasma IL-6 concentrations were measured as described previously (*27*). All analyses were performed in R (version 4.1.3). From the 2516-participant BIOSTAT-CHF index cohort, patients were selected based on ischemic HF etiology, available peripheral edema status, and absence of auscultatory signs of pulmonary congestion, yielding a final analytic cohort of 418 patients. The independent associations of peripheral edema presence and LVEF (categorized as >30% vs. ≤30%) with plasma IL-6 concentrations was assessed by linear regression. Statistical significance was defined as $P<0.05$.

Invasive hemodynamic measurements

Pressure-volume measurements of the RV were performed in a closed-chest protocol under echocardiographic guidance. A 1.0-F pressure conductance catheter (PVR-1035, Millar) was pre-soaked in 0,9% NaCl for at least 30 minutes and subsequently calibrated for pressure. After induction of anesthesia using 1.5-2% isoflurane, the catheter was inserted into the RV via a 22-gauge cannula. Intra-ventricular catheter position was optimized under echocardiographic guidance. Body temperature was assessed using a rectal temperature probe and maintained within a stable range of 36 to 37 °C. Parallel conductance was measured by injecting hypertonic (10%) saline solution into the right jugular vein. The conversion from conductance to volume was applied by placing the catheter into known volumes of the calibration cuvette, filled with heparinized blood. All analyses were performed in LabChart 8 (ADInstruments, New Zealand).

Immunofluorescence staining and analysis

Cryosection slides were brought to room temperature and fixed in 4% paraformaldehyde for 20 minutes. After rehydration in PBS, sections were permeabilized with 0.5 % Tween 20 for 25 minutes and blocked with 5% bovine serum albumin (BSA) and 0.1% Tween 20 for 1 hour at room temperature. Samples were then circled using a hydrophobic barrier pen. Primary antibodies diluted in 1% BSA were applied, and sections were incubated overnight at 4 ºC in a humified chamber. The following primary antibodies were used: anti-CD68 (abcam, AB53444, clone FA-11) at a 1:200 dilution, anti-PDGFRA (abcam, AB203491, clone EPR22059-270) at a 1:400 dilution, and anti-CD31 (R&D Systems, AF3628, polyclonal) at a 1:400 dilution.

On the following day, sections were washed and stained for 1 hour at room temperature with the corresponding secondary antibodies, together with labeled wheat germ agglutinin (Invitrogen, W11261). The following secondary antibodies were used: donkey anti-goat IgG, Alexa Fluor 555 (Invitrogen, A21432), donkey anti-rat IgG, Alexa Fluor Plus 647 (Invitrogen, A48272), and donkey anti-rabbit IgG, Alexa Fluor 750 (abcam, AB175728). After washing, sections were stained with 300 nM DAPI (Thermo Fisher Scientific, D1306) for 5 minutes and washed again. Sections were finally covered in mounting medium (Fluoromount-G, Southern Biotech) and stored light-protected at room temperature. Imaging was performed using a slider scanner (Axioscan 7, Zeiss). Analysis was performed using QuPath software (QuPath v.0.6.0/ v.0.7.0). Whole-section images were manually annotated into the following anatomical regions of interest: LV free wall,

LV septal wall, RV, right atrium, and left atrium. Nuclear segmentation was performed using the QuPath cell detection tool based on DAPI signal, with cell boundaries estimated by expansion of the nuclear mask. Fluorescence intensity thresholds for CD68 and PDGFRA classification were determined empirically based on secondary-antibody-only control stainings and an CD31+ endothelial cell negative reference population. Cells with mean intensity exceeding the run-specific CD68 threshold were classified as CD68+ myeloid cells; cells with mean intensity exceeding the run-specific PDGFRA threshold were classified as fibroblasts.

Untargeted metabolomics of murine plasma samples

Blood was collected into labeled EDTA-coated tubes and centrifugated (4°C, 2000 x g, 10 minutes). Plasma was stored at −80°C until analysis. After thawing on ice, 50 µL aliquots of plasma samples were extracted by addition of 200µL MeOH (including internal standards). Samples were then thoroughly vortexed for 10 minutes at 4 °C and subsequently incubated at -20 °C for 20 minutes. Ultimately, the extracted samples were centrifuged for 10 minutes at 15,000g and 4 °C, and 100 µL of the supernatants were transferred to analytical glass vials for LC-MS/MS analysis. LC-MS/MS analysis was performed on a Vanquish Horizon UHPLC system coupled to an Orbitrap Exploris 240 high-resolution mass spectrometer (Thermo Scientific, MA, USA) in negative and positive ESI (electrospray ionization) mode. Chromatographic separation was carried out on an Atlantis Premier BEH Z-HILIC column (Waters, MA, USA; 2.1 mm x 100 mm, 1.7 µm) at a flow rate of 0.25 mL/min. The mobile phase consisted of water:acetonitrile (9:1, v/v; mobile phase A) and acetonitrile:water (9:1, v/v; mobile phase B), which were modified with a total buffer concentration of 10 mM ammonium acetate (negative mode) and 10 mM ammonium formate (positive mode), respectively. The aqueous portion of each mobile phase was pH-adjusted. Column temperature was maintained at 40°C, the autosampler was set to 4°C and sample injection volume was 5 µL. Analytes were recorded via a full scan with a mass resolving power of 120,000 over a mass range from 60 – 900 m/z. To obtain MS/MS fragment spectra, data-dependent acquisition was carried out. All experimental samples were measured in a randomized manner. Pooled quality control (QC) samples were prepared by mixing equal aliquots from each processed sample. For determination of background signals and subsequent background subtraction, an additional processed blank sample was recorded. Data was processed using MS DIAL 4.9.221218. Level 1 feature identification was based on an in-house library for metabolomics (EMBL-MCF 2.0) using accurate mass, isotope pattern, MS/MS fragmentation, and retention time information and a minimum matching score of 80%.

FITC dextran measurement

Mice were fasted for 4–6 h, then orally gavaged with 150 µL of 80 mg/mL FITC-dextran (4 kDa, Sigma-Aldrich FD4) prepared in sterile PBS. Blood was collected via facial vein puncture 4 h post-gavage using MiniCollect CAT Serum Separator tubes (Greiner Bio-One, Austria, cat. no. 450533) and centrifuged to isolate serum. Serum samples were diluted in PBS, loaded onto 96-well plates, and fluorescence was measured at 530 nm following excitation at 485 nm. Intestinal permeability was expressed as relative fluorescence units after subtraction of PBS background fluorescence.

Flow cytometry of murine samples

Peripheral blood was collected by facial vein puncture in EDTA-coated tubes, and erythrocytes were lysed in RBC lysis buffer (Miltenyi Biotec). Spleens and bone marrow were harvested after cervical dislocation. Single-cell suspensions of bone marrow were obtained by flushing 1 dissected femur with 2 mL ice-cold PBS. Spleens were mechanically dissociated by passing tissue through 40 µm cell strainers and erythrocytes were lysed in RBC lysis buffer for 2 minutes (Miltenyi Biotec). For flow cytometry, cells were incubated with fluorophore-conjugated monoclonal antibodies (1:200 dilution) in FACS buffer (2 mM EDTA, 2% FCS in PBS) for 30 minutes on ice. The following antibodies were used: anti-Ly6G PE (BD Bioscience, 561104, clone 1A8), anti-CD45 FITC (BD Bioscience, 553080, clone 30-F11), anti-CD11b APC-Cy7 (BD Bioscience, 557657, clone M1/70), anti-CD31 PE-Cy7 (BioLegend, 160213, clone W18222B), anti-CD117 PE-Cy7 (BD Bioscience, 558163, clone 2B8), anti-Ly6C APC (BD Bioscience, 560595, clone AL-21), anti-CD18 FITC (BioLegend, 101405, clone M18/2), anti-IgE BV421 (BD Bioscience, 564207, clone R35-72), anti-CD45 PerCP-Cy5.5 (BD Bioscience, 550994, clone 30-F11), anti-Ter119 PE (BD Biosciences, 553673, clone TER-119), anti-CD3e PE (BD Biosciences, 553064, clone 145-2C11), anti-B220 PE (BioLegend, 103208, clone RA3-6B2), anti-Ly6C FITC (BD Bioscience, 553104, clone AL-21), anti-F4/80 PE-Cy7 (eBioscience, 25-4801-82, clone BM8), anti-B220 APC (BD Biosciences, 553092, clone RA3-6B2), anti-CD4 PE-Cy7 (BD Biosciences, 552775, clone RM4-5), anti-CD8b BV510 (BD Biosciences, 740155, clone H35-17.2), anti-CD11b PE (BD Pharmingen, 557397, clone M1/70), anti-GR-1 PE (BioLegend, 108408, clone RB6-8C5), anti-c-Kit PE-Cy7 (BioLegend, 105814, clone 2B8), anti-CD150 APC (BioLegend, 115910, clone TC15-12F12.2), anti-Sca-1 PerCP-Cy5.5 (BioLegend, 122524, clone E13-161.7), anti-CD16/32 APC-Cy7 (BioLegend, 101328, clone 93), and anti-CD34 BV421 (BD Biosciences, 562608, clone RAM34). For intracellular IL-6 staining, the BD Cytofix/Cytoperm kit (BD Biosciences, #554714) was used according to the manufacturer's instruction following surface staining. Fixed and permeabilized cells were staining with anti-IL-6 APC (BioLegend, 504507, clone MP5-20F3) at a 1:100 dilution for 30 minutes on ice. Data acquisition was performed using a FACSVerse cell analyzer (BD Biosciences). The gating strategy used for flow cytometric analysis was as follows: All cells were pre-gated on single cells (FSC-A vs. SSC-A, and SSC-W vs. SSC-H). Bone marrow LSK were identified as Lin- (Ter119, CD3, B220, CD11b, GR-1) c-kit+ Sca-1+ cells. These were further divided into long-term hematopoietic stem cells (HSCs; Lin- c-kit+ Sca-1+ CD150+). Granulocyte macrophage progenitors (GMP) were identified as Lin- c-kit+ Sca-1- CD16/32+ CD34+. Marrow monocytes were identified as CD45+ CD11b+ Ly6G-, marrow neutrophils as CD45+ CD11b+ Ly6G+ cells. Blood and spleen leukocytes were identified as CD45+ singlets. Myeloid cells were further subdivided into CD11b+Ly6G+ cells (neutrophils), CD11b+ Ly6G- Ly6C-high cells (Ly6C-high monocytes) and CD11b+ Ly6G- Ly6C-low cells (Ly6C-low monocytes). Lymphoid cells were further subdivided into $B220^+$ cells (B cells) and $CD3^+$ cells (T cells), with T cells further gated based on CD4+ and CD8+ expression. Data were analyzed using FlowJo software (BD Biosciences, v.10.9.0).

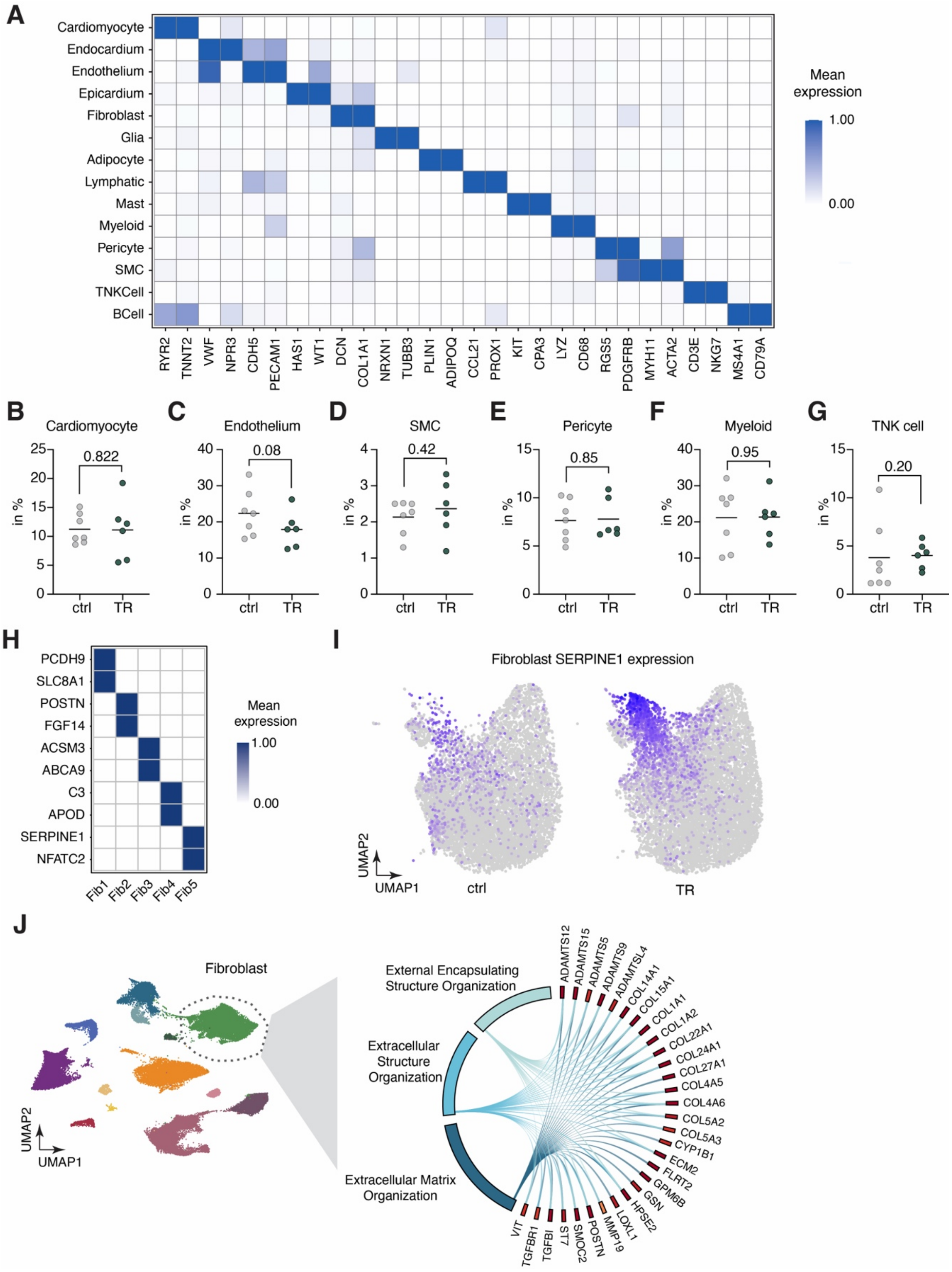
A
Cardiomyocyte
Endocardium
Endothelium
Epicardium
Fibroblast
Glia
Adipocyte
Lymphatic
Mast
Myeloid
Pericyte
SMC
TNKCell
BCell
RYR2 TNNT2 VWF NPR3 CDH5 PECAM1 HAS1 WT1 DCN COL1A1 NRXN1 TUBB3 PLIN1 ADIPOQ CCL21 PROX1 KIT CPA3 LYZ CD68 RGS5 PDGFRB MYH11 ACTA2 CD3E NKG7 MS4A1 CD79A
Mean expression
1.00
0.00
B Cardiomyocyte 0.822
C Endothelium 0.08
D SMC 0.42
E Pericyte 0.85
F Myeloid 0.95
G TNK cell 0.20
in %
ctrl TR
H
PCDH9
SLC8A1
POSTN
FGF14
ACSM3
ABCA9
C3
APOD
SERPINE1
NFATC2
Fib1 Fib2 Fib3 Fib4 Fib5
Mean expression
1.00
0.00
I
Fibroblast SERPINE1 expression
UMAP2
UMAP1
ctrl
TR
J
Fibroblast
UMAP2
UMAP1
External Encapsulating Structure Organization
Extracellular Structure Organization
Extracellular Matrix Organization
ADAMTS12 ADAMTS15 ADAMTS5 ADAMTS9 ADAMTSL4 COL14A1 COL15A1 COL1A1 COL1A2 COL22A1 COL24A1 COL27A1 COL4A5 COL4A6 COL5A2 COL5A3 CYP1B1 ECM2 FLRT2 GPM6B GSN HPSE2 LOXL1 MMP19 POSTN SMOC2 ST7 TGFBI TGFBR1 VIT

**Fig. S1. Single-nucleus RNA sequencing of human left ventricular tissue in human TR.**

(**A**) Heatmap of mean gene expression for cell-type marker genes across 13 annotated cell populations identified by snRNA-seq of human LV tissue. Color scale indicates mean expression. (**B-G**) Proportions (% of total nuclei) of indicated cell types in control and TR patients. Data show means with individual data points. *P*-values were calculated using WLS regression. (**H**) Heatmap of mean marker gene expression across LV fibroblast subclusters. (**I**) UMAP plots of LV fibroblast subclusters showing SERPINE1 expression in ctrl (left) and TR (right) samples. (**J**) Chord diagram depicting fibroblast-enriched gene ontology terms and associated top contributing genes.

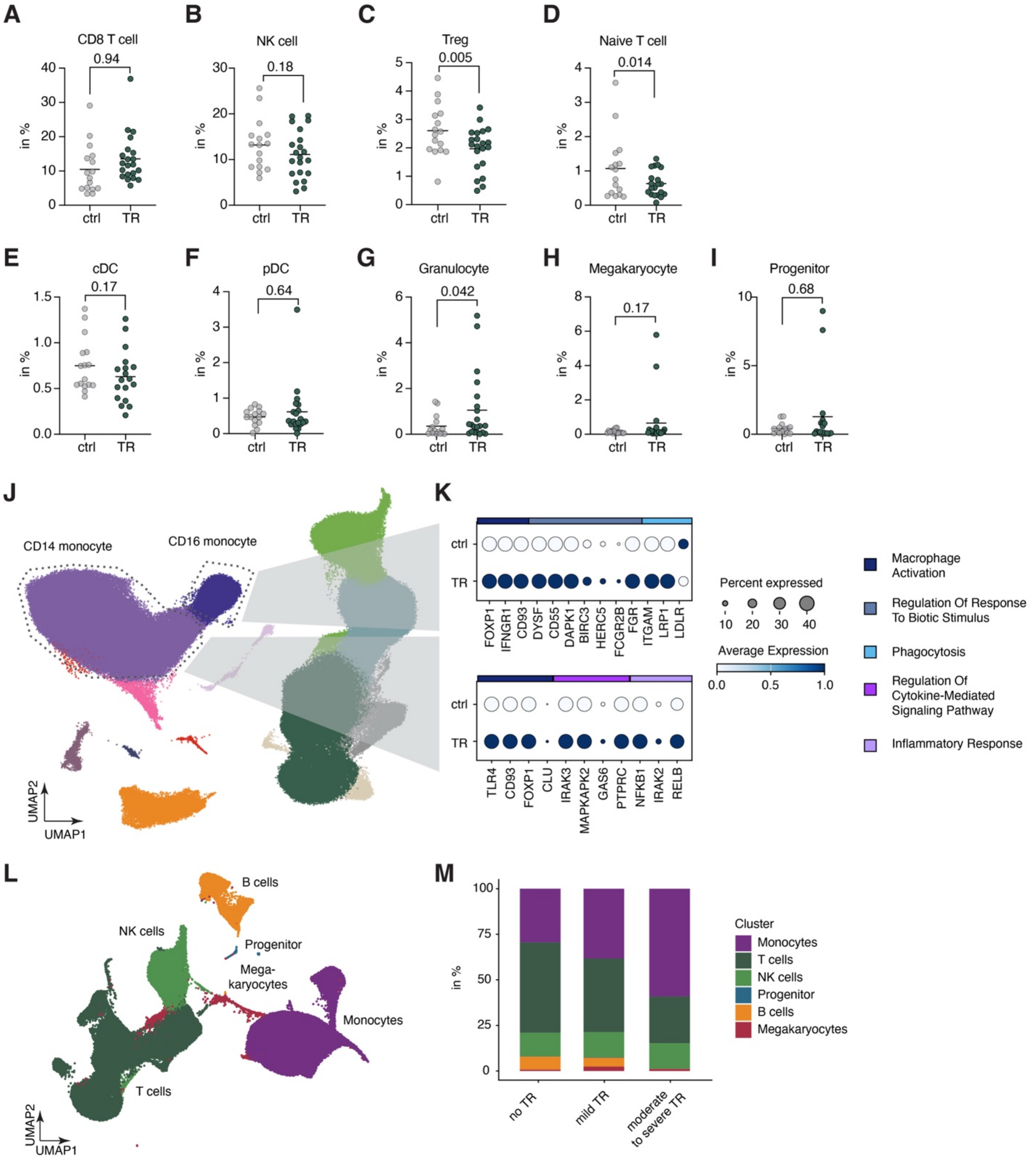


**Fig. S2. Immune cell composition and monocyte transcriptional activation in blood of TR patients.**

(**A-I**) Proportions (in % of total PBMCs) of circulating immune cell subsets in ctrl (gray) and TR (green) patients. Data show means with individual data points. *P*-values were calculated using WLS regression with Benjamini-Hochberg correction. (**J-K**) Enrichment of gene sets grouped by GO biological process annotation. Dot plots showing expression of select marker genes in CD16+ monocytes (top) and CD14+ monocytes (bottom) comparing control (ctrl) and TR conditions. Dot size encodes percent expressed; color intensity encodes average expression. Associated gene ontology terms shown at right. (**L**) UMAP of PBMCs colored by annotated cluster profiled in an additional cohort from Kneuer et al. (*16*). (**M**) Stacked bar charts showing proportional composition of blood immune cell clusters across TR severity groups.

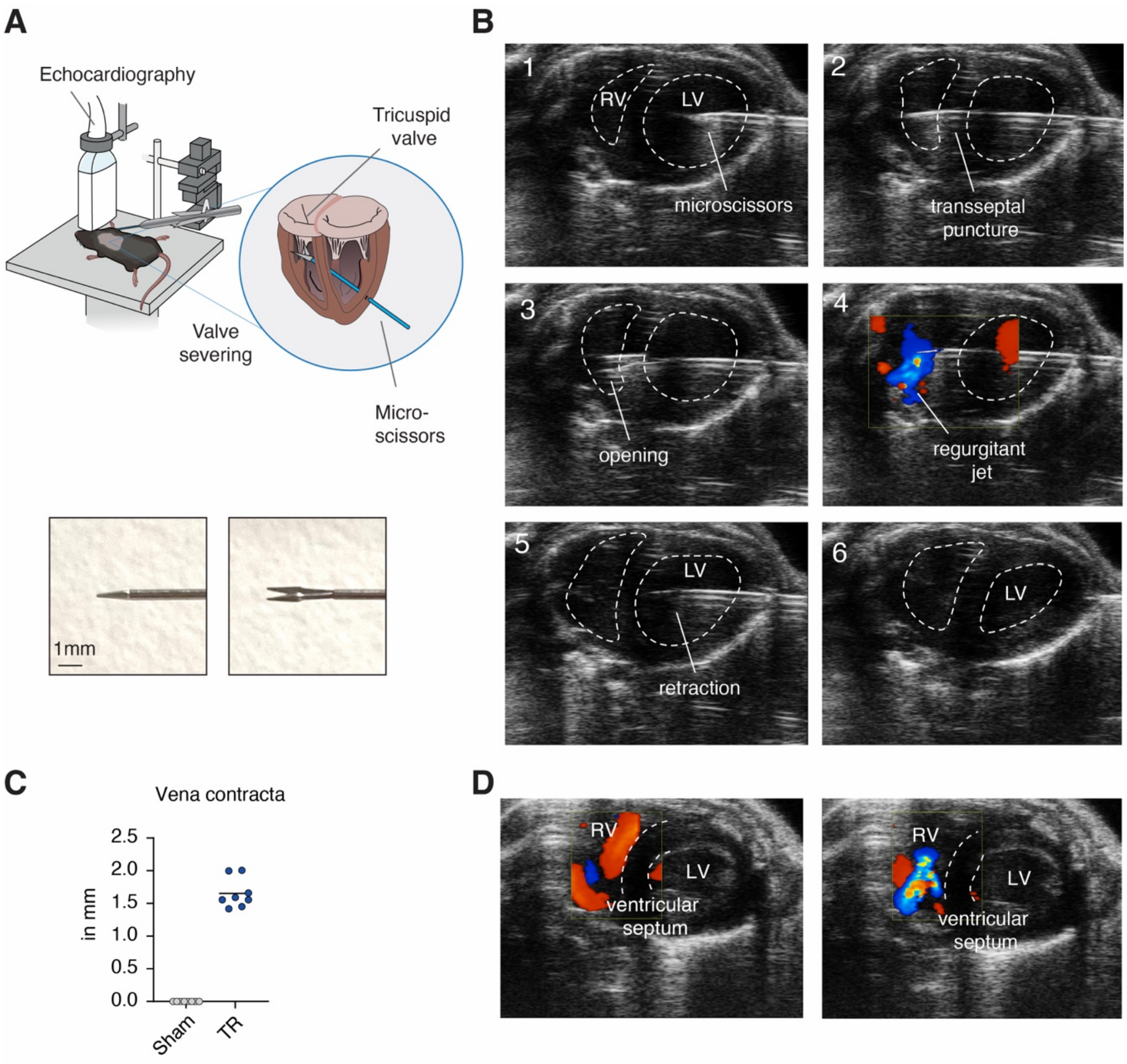


**Fig. S3. Minimally invasive induction of TR in mice.**

(**A**) Schematic of the experimental preparation: the tricuspid valve is accessed and severed using microscissors under echocardiographic guidance (top). Photographs (bottom) show the microscissors in closed (left) and open (right) configurations. Scale bar, 1 mm. (**B**) Sequential echocardiographic images (frames 1–6) documenting key procedural steps: insertion of microscissors (1), transseptal puncture (2), leaflet opening (3), appearance of regurgitant jet by color Doppler (4), instrument retraction (5), and post-procedural view (6). RV, right ventricle; LV, left ventricle. (**C**) Quantification of vena contracta width (mm) in sham and TR mice at 28 days post-intervention (mean with individual data points). (**D**) Representative color Doppler echocardiographic images during diastole (left) and systole (right) confirming absence of interventricular shunting in TR mice.

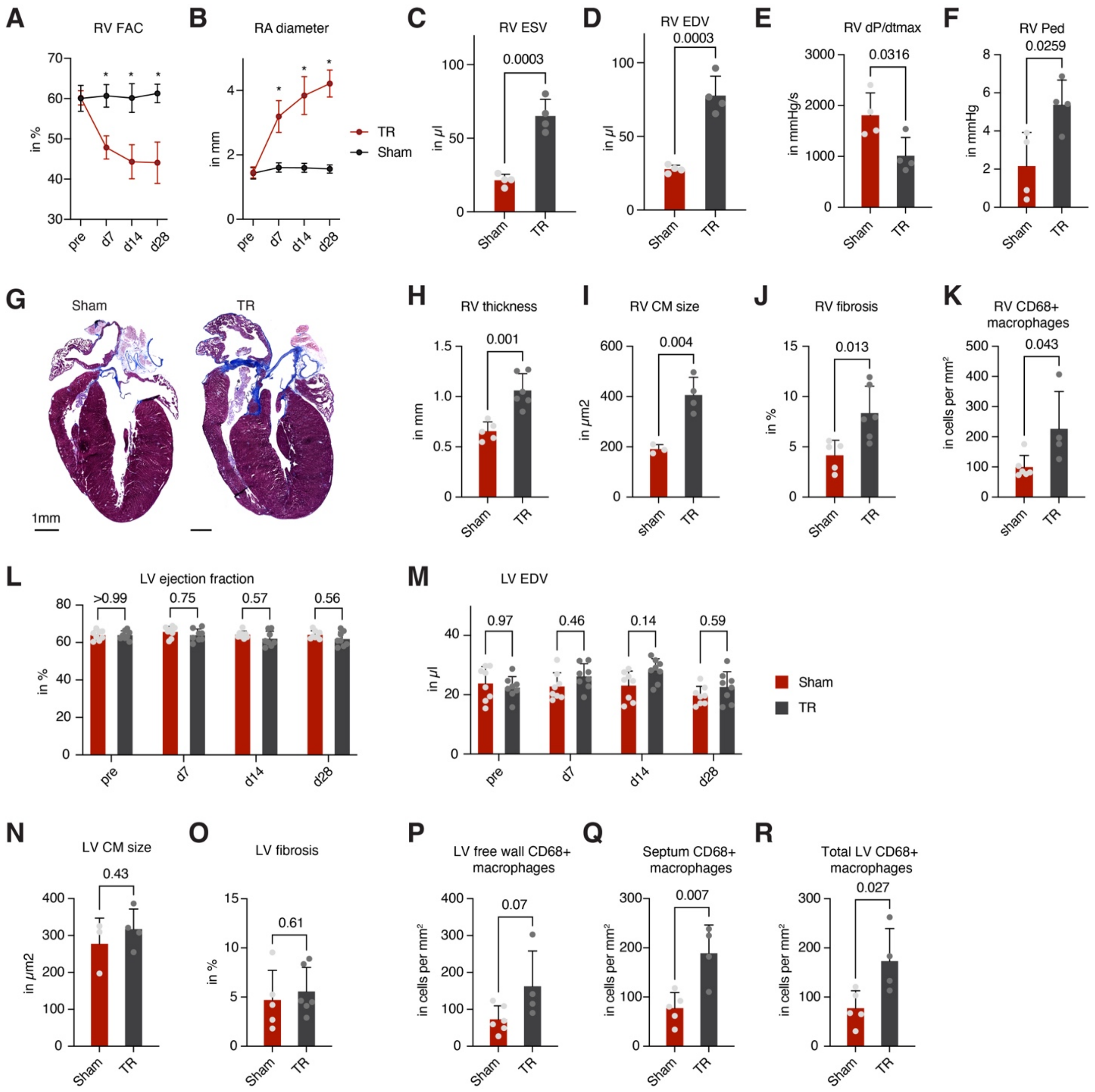


**Fig. S4. Cardiac remodeling following isolated TR.**

(**A**) Longitudinal echocardiographic characterization of RV function (RV fractional area change, in %) in the mouse TR model (n = 8, two-way ANOVA with Sidak's multiple comparison, mean ± SD). *$P$ < 0.05. (**B**) Serial echocardiographic quantification of RA diameter over time (n = 8, two-way ANOVA with Sidak's multiple comparison mean ± SD). *$P$ < 0.05. (**C-F**) Hemodynamic parameters from invasive RV pressure-volume catheterization at d28, comparing sham and TR mice (n = 4, unpaired t-test, mean ± SD). (**G**) Representative Masson's trichrome-stained long axis-sections of sham and TR hearts. Scale bar, 1mm. (**H**) RV free wall thickness at d28, reflecting hypertrophic remodeling in response to chronic volume overload (n = 4-5, unpaired t-test, mean ± SD). **(I)** RV cardiomyocyte cross-sectional area at d28 based on histological evaluation (n = 3-4, unpaired t-test, mean ± SD). **(J)** RV interstitial fibrosis quantified as collagen-positive area fraction (%) from Masson's trichrome-stained sections at d28 (n = 5-6, unpaired t-test, mean ± SD). **(K)** RV CD68+ macrophage density, quantified by immunofluorescence staining at d28 post-TR (n = 4-6, unpaired t-test, mean ± SD). (**L-M**) LV ejection fraction (in %) and LV EDV (in µl) across timepoints (n = 8, two-way ANOVA with Sidak's multiple comparison mean ± SD). (**N**) LV cardiomyocyte cross-sectional area (n = 3-5, unpaired t-test, mean ± SD). (**O**) LV interstitial fibrosis (in % of whole annotated tissue) based on Masson's trichrome staining (n = 5-6, unpaired t-test, mean ± SD). (**P–R**) CD68+ macrophage density in LV free wall, septum, and total (n = 4-5, unpaired t-test, mean ± SD).

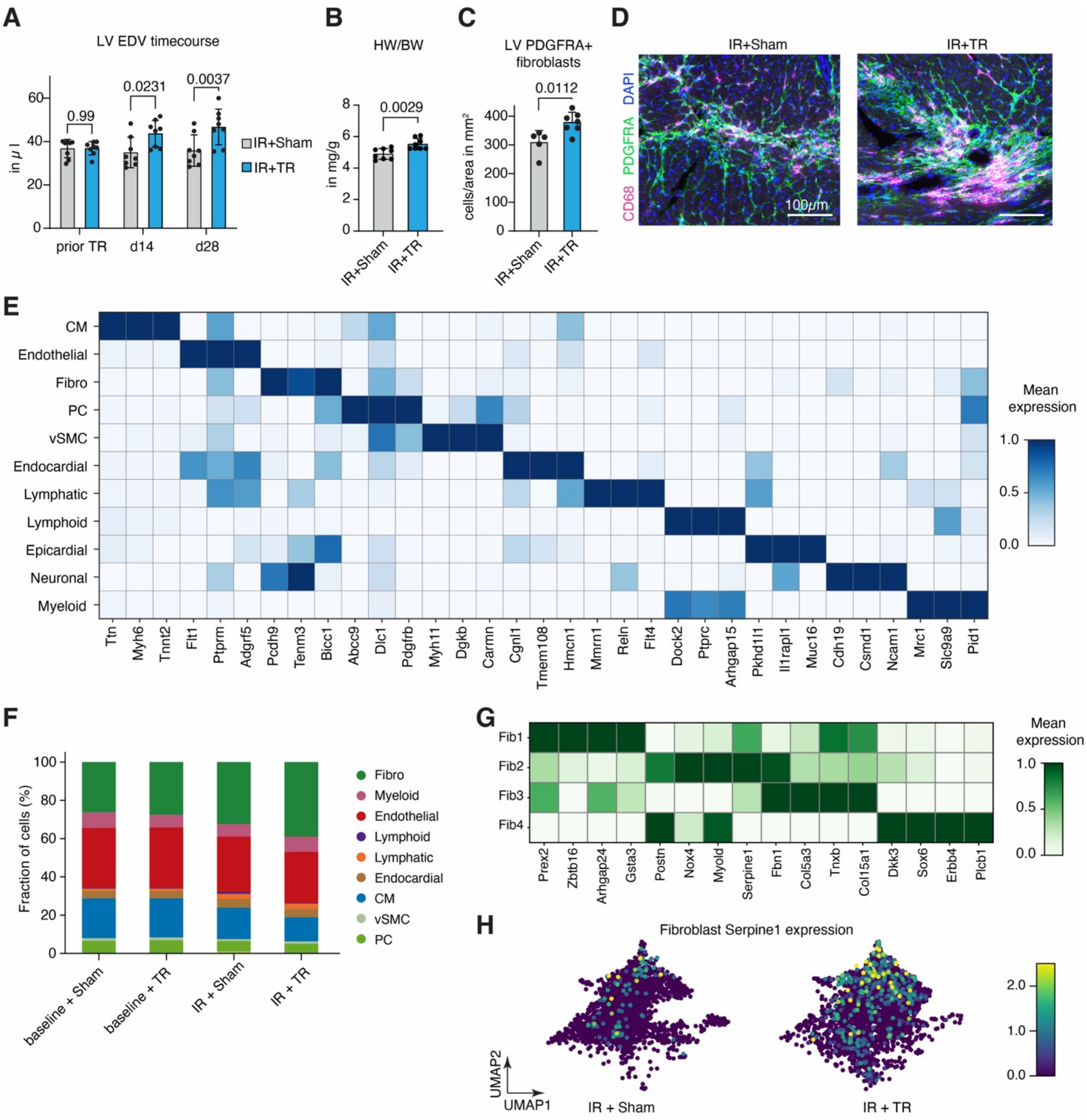


**Fig. S5. LV remodeling in mice with ischemic cardiomyopathy and superimposed TR.**
(**A**) LV end-diastolic volume (EDV) timecourse at baseline (prior to TR), d14, and d28 in mice with IR injury and superimposed sham (IR+sham, gray) or TR induction (IR+TR, blue; n = 8, two-way ANOVA with Sidak's multiple comparison, mean ± SD). (**B**) Heart-to-body weight ratio (HW/BW) at d28 (n = 8, unpaired t-test, mean ± SD). (**C**) PDGFRA+ LV fibroblast density (in cells/mm²) in IR+sham versus IR+TR (n = 5-7, unpaired t-test, mean ± SD). (**D**) Representative immunofluorescence images of LV sections stained for PDGFRA (green), CD68 (magenta), and DAPI (blue) in IR+sham and IR+TR. (**E**) Heatmap of mean marker gene expression across 11 annotated LV cell populations from snRNA-seq. (**F**) Stacked bar plots showing proportional composition of major cardiac cell types across all condition based on snRNA-seq. (**G**) Heatmap

of mean marker gene expression across LV fibroblast subclusters. (**H**) UMAP plots of LV fibroblast subclusters showing Serpine1 expression in IR+sham (left) and IR+TR (right) samples.

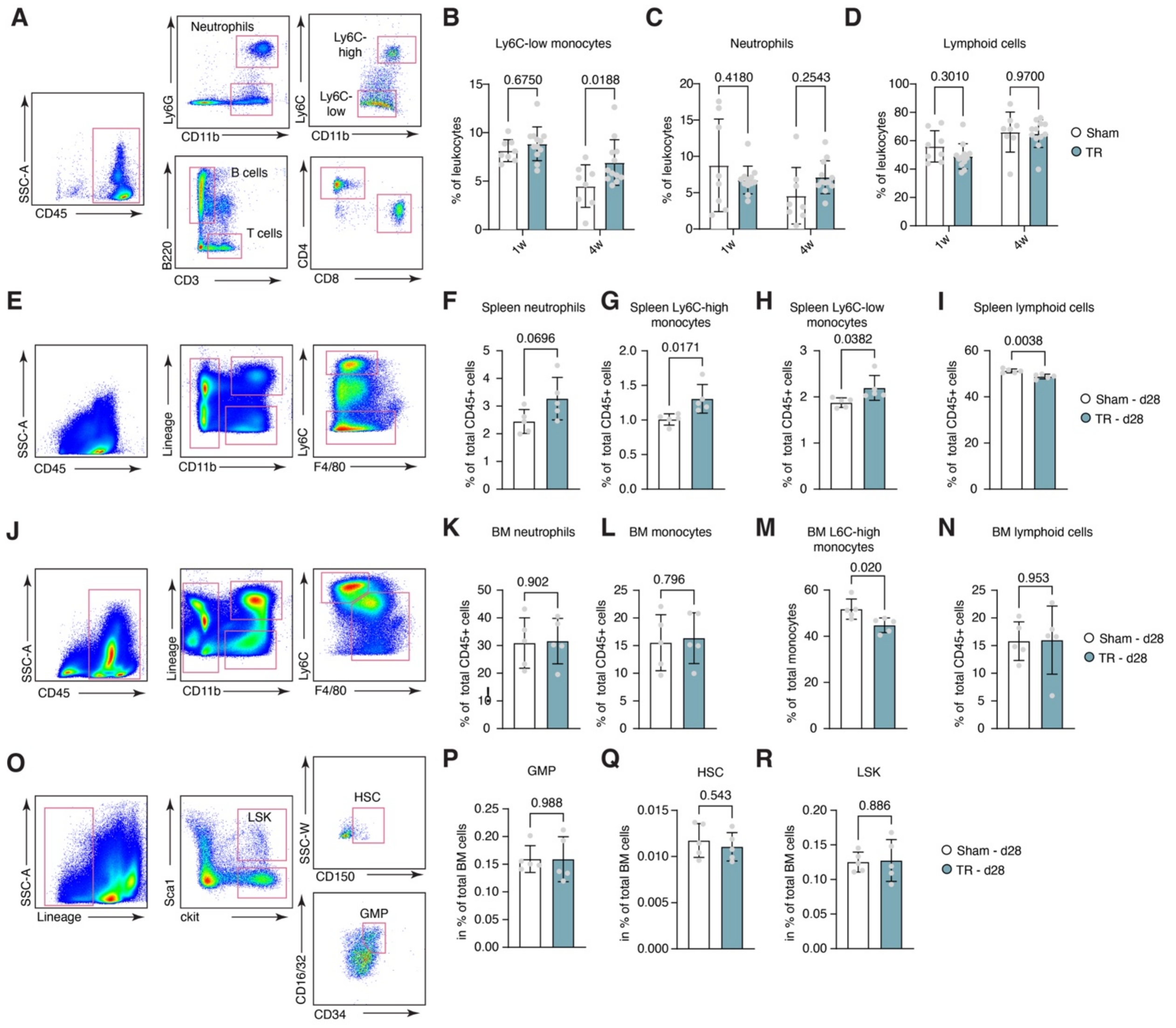


**Fig. S6. Flow cytometric analysis of leukocytes in blood, spleen, and bone marrow in TR.**

**(A)** Gating strategy for blood leukocytes. (**B–D**) Blood Ly6C-low monocytes, neutrophils, and lymphoid cells (% leukocytes) at 1w and 4w post-TR (n = 8-12, two-way ANOVA with Sidak's multiple comparison, mean ± SD). **(E)** Gating strategy for splenic leukocytes. (**F–I**) Relative percentage (in % CD45+ cells) of spleen neutrophils, Ly6C-high monocytes, Ly6C-low monocytes, and lymphoid cells (n = 5, unpaired t-test, mean ± SD). **(J)** Gating strategy for bone marrow leukocytes. (**K–N**) Relative frequency of bone marrow neutrophils, total monocytes, Ly6C-hi percentage of total monocytes, and lymphoid cells (n = 5, unpaired t-test, mean ± SD). **(O)** Gating strategy for bone marrow stem cells and progenitor cells. (**P–R**) Percentage (in % of total bone marrow cells) of HSC, LSK, and GMP populations (n = 5, unpaired t-test, mean ± SD).

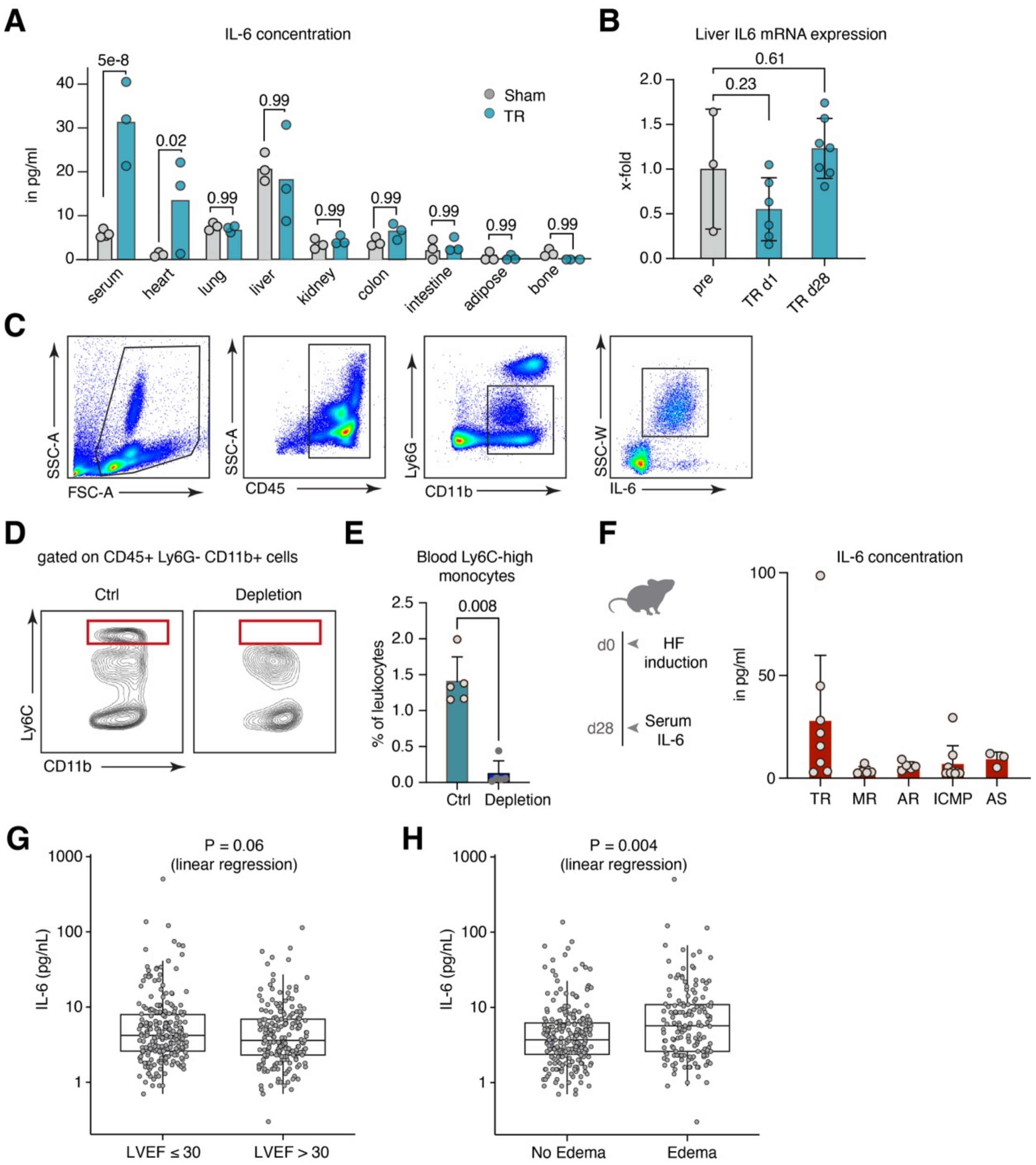


**Fig. S7. IL-6 tissue distribution, monocyte-dependent secretion, and clinical associations in heart failure.**

(**A**) IL-6 protein concentration (in pg/ml) measured across tissues (serum, heart, lung, liver, kidney, colon, intestine, adipose, bone) in sham (gray) and TR (blue) mice 28 days after intervention (n = 3-4, two-way ANOVA with Sidak's multiple comparison, mean with individual data points). (**B**) Hepatic IL-6 mRNA expression at d1 and d28 post-TR relative (in x-fold compared to sham, n = 3-7, one-way ANOVA with Tukey's multiple comparison, mean ± SD). (**C**) Flow cytometry gating strategy for intracellular IL-6 detection in blood monocytes. (**D**) Representative flow cytometry plots of CD45+ Ly6G- CD11b+ cells from control and monocyte-depleted mice. (**E**) Frequency of blood Ly6C-high monocytes (% leukocytes) in control vs.

depleted mice (n = 5, unpaired t-test, mean ± SD). (**F**) Serum IL-6 (in pg/ml) across murine HF entities: TR, mitral regurgitation (MR), aortic regurgitation (AR), ischemic cardiomyopathy (ICMP), and aortic stenosis (AS, induced by transverse aortic constriction). Blood was sampled on d28 after disease induction. (**G-H**) Serum IL-6 (in pg/nl, log scale) stratified by LV EF <30% vs. ≥30% (G) or by presence or absence of peripheral edema (H) in the BIOSTAT-CHF cohort (*27*). *P*-values were calculated by linear regression.

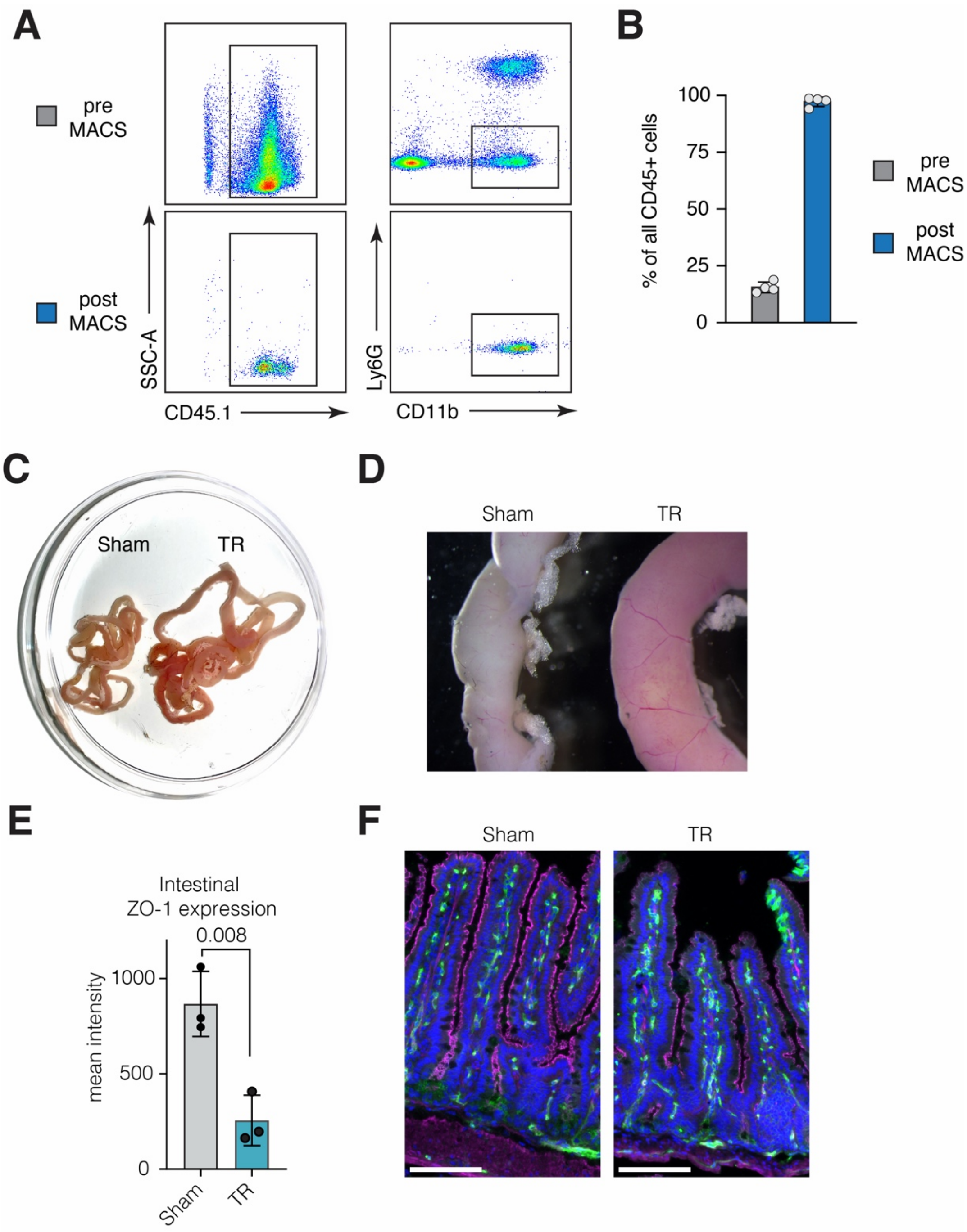


**Fig. S8. Validation of monocyte isolation protocol and assessment of gut barrier function.**
(**A**) Representative flow cytometry plots showing CD45.1 vs. SSC-A (left) and Ly6G vs. CD11b (right) of blood cells before (pre-MACS, top) and after (post-MACS, bottom) magnetic-activated cell sorting. (**B**) Quantification of monocyte enrichment as percentage of total CD45$^+$ cells pre- and post-MACS. (**C**) Gross photograph of intestinal segments from sham and TR mice at d28. (**D**) Stereomicroscopic photographs of intestinal cross-sections from Sham and TR mice at d28. (**E**) Intestinal ZO-1 protein expression (mean fluorescence intensity) based on immunofluorescence

staining in sham vs. TR mice (n = 3, unpaired t-test, mean ± SD). (**F**) Representative immunofluorescence images of intestinal sections stained for ZO-1 (magenta), CD31 (green) and DAPI (blue) in sham and TR.

**Table S1.**
Clinical characteristics of the retrospective propensity score–matched cohort.

**Table S2.**
Clinical and demographic characteristics of patients included in PBMC snRNA-seq analysis.

**Table S3.**
Patient-level metadata for the plasma proteomics cohort comparing TR patients to controls.

**Table S4.**
Clinical metadata and IL-6 measurements in patients before and after TTVR.

**Movie S1. Echocardiography-guided induction of tricuspid regurgitation in mice.**
Real-time echocardiographic recording of tricuspid valve severing using microscissors via trans-LV and transseptal access. Tricuspid regurgitation is confirmed by color Doppler imaging.
